# Intermediate Evolutionary State of Motile Sperm and Pollen Tubes in the Extant Gymnosperm *Cycas revoluta*

**DOI:** 10.1101/2025.03.22.644562

**Authors:** Yukiho Toyama, Satohiro Okuda, Takamasa Suzuki, Tetsuya Higashiyama

**Author notes:** Yukiho Toyama; The University of Tokyo, 7-3-1 Hongo, Bunkyo, Tokyo, 113-0033, Japan.

## Abstract

The transformation of male reproductive cells is a significant event in the evolution of land plant fertilization systems from zooidogamy to siphonogamy. Basal plants such as bryophytes and pteridophytes have motile sperm, whereas most seed plants have non-motile sperm, delivered by a pollen tube. Despite being seed plants, gymnosperm cycads and ginkgo uniquely form highly multi-flagellated and large motile sperm within pollen tubes. However, the evolutionary state of these male reproductive cells remains unknown. We clarified the gene expression profiles of *Cycas revoluta* pollen tubes and motile sperm that swam toward female reproductive cells. Male cycad cells had fewer genes associated with transcription, translation, and related processes, which is consistent across land plants. Similar to other male gametes, a sperm-specific histone variant may contribute to transcriptional regulation via chromatin condensation. Plasma membrane-localized proteins probably involving male–female interactions revealed that cycad sperm express both factors found in motile plant sperm and seed plant sperm cells, suggesting a transitional state from motile to non-motile male gametes. In contrast, cycad pollen tubes lacked plasma membrane-localized receptor genes previously reported in angiosperms and cell wall-associated factors involved in pollen tube tip growth. These results support the hypothesis that the function of the cycad pollen tube is confined to the haustoria as previously reported. These results indicate a molecular intermediate state of the cycad fertilization mechanism with motile sperm and pollen tubes, providing important insights into the evolution of land plant fertilization.

**Significant Statement:** Cycads and ginkgo, which possess primitive motile sperm and derived pollen tubes, are important for understanding the evolution of land plant fertilization from zooidogamy to siphonogamy. We explored gene expression in living, capacitated motile sperm and pollen tubes of *Cycas revoluta* in a tissue-specific transcriptome analysis. Similar to other land plants, the transcriptional repression mechanism has been suggested in these male cells. Plasma membrane-localized genes revealed the characteristics of motile and non-motile sperm, while molecular evidence of pollen tube guidance and tip growth, as seen in angiosperms, was lacking in cycads. A molecular intermediate state of the cycad fertilization mechanism was clarified, providing important insights into the evolutionary process of land plant fertilization.

## Introduction

The sex fertilization system in which male and female gametes carrying genetic information fuse together is an essential reproductive process. During land plant evolution, the fertilization system transited from zooidogamy (“motile sperm”) to siphonogamy (“pollen tube-mediated”). This transition is represented by a large-scale change in the morphology of male reproductive cells, notably the loss of sperm motility and the acquisition of pollen tubes extending from pollen (1, 2). Basal land plants such as bryophytes and pteridophytes have flagellated, motile sperm produced in the antheridium that swim directly to the egg cell. In contrast, most seed plants, including angiosperms and some gymnosperms such as conifers and gnetophytes have non-motile sperm without flagella within the pollen tube, which depend on the pollen tube for movement to the egg cell (3). The motile sperm of basal land plants and non-motile sperm cells of angiosperms share a common mechanism, in which nuclear chromatin is highly condensed and transcriptional activity is suppressed, as in animal sperm (1, 4). This mechanism relies on the replacement of somatic histones in sperm with sperm-specific nuclear basic proteins (SNBPs). Bryophytes contain protamine-like SNBPs, whereas pteridophytes have both protamine and histone types and angiosperms have only histone types. All of these SNBP types promote nuclear chromatin condensation (5). Based on these similar gene expression dynamics, upstream transcription factors controlling the development and differentiation of male gametes, as well as fertilization-related gamete fusion genes, are shared by both types of male gametes (6–10).

However, there are significant differences in the processes that occur before gamete fusion. Although a plasma membrane-localized receptor protein has been identified as a factor related to sperm chemotaxis in bryophytes, few studies have examined female signal reception in non-motile angiosperm sperm cells (11). Conversely, functional analyses of angiosperm pollen tubes are common; this structure was acquired after the emergence of seed plants. Angiosperm pollen tubes are characterized by tip growth a short time after pollination and a molecular mechanism for receiving female signals for guidance to egg cells (12). Several specifically expressed membrane-localized receptor-like kinases (RLKs) have been identified (13). Due to the specificity of angiosperm-like pollen tubes, it is difficult to determine how such molecular mechanisms were acquired. Thus, although both types of male reproductive cells partially retain their basic molecular mechanisms, differences in motility make it difficult to directly compare and trace the evolutionary trajectories of molecular mechanisms involved in the processes preceding gamete fusion between basal land plants and angiosperms.

This study focused on cycads and ginkgo, which are the only extant zooidogamous gymnosperm seed plants (14). Their motile sperm are generated in pollen tubes and reach the egg by swimming. These plants exhibit a significant evolutionary state that combines characteristics of zooidogamy and siphonogamy, containing essential information for inferring the transition of male reproductive cells during evolution. Despite this evolutionary significance, the features are morphologically unique compared to other plants. Their sperm are large and have extremely large numbers of flagella, ranging from several thousand to tens of thousands, suggesting a lack of nuclear condensation (14–16). Based on their morphological characteristics, cycad pollen tubes have no tip growth mechanism and function as haustoria by consuming metabolites in the nucellar tissue of the ovules to nurture the developing sperm (17). However, little is known about the gene expression dynamics of the unique reproductive cells of cycads and ginkgo, and their molecular evolutionary state remains completely unknown.

This lack of data is caused by the difficulty of sampling, as cycads and ginkgo are slow-growing plants that can take several years to several decades to fruit (18, 19). Reproductive tissues can be collected only once per year during the fertilization period, which lasts just a few days (20). Continuous observations throughout the fertilization season have shown that *Cycas revoluta*, which is native to Japan, exhibits an intermittent but extended fertilization period of several months in northern to southern Japan. In this study, we utilized *C. revoluta* to explore the gene expression profiles of cycad motile sperm and pollen tubes based on tissue-specific transcriptome analysis using isolated intact sperm and other reproductive and non-reproductive tissues. Despite the unique morphological characteristics of cycad male reproductive cells, a common transcriptional repression mechanism using an estimated SNBP was suggested. Furthermore, a search for factors predicted to be localized in the cell membrane of cycad sperm and pollen tubes allowed us to estimate how molecules are shared with other land plants. This study is the first to capture the molecular dynamics of this fertilization system combining motile sperm and pollen tubes, and provides evidence for the reconsideration of the evolutionary state of the fertilization stage.

## Results

### Isolation of motile and fertile cycad sperm

Four types of male and female reproductive tissues and three non-reproductive tissues were collected for RNA sequencing. About 3 months after pollination, the female reproductive tissue matured within the ovule, followed by growth of the pollen tube. Mature reproductive cells, including pollen tubes, neck cells, and egg cells were observed in the inner region of the upper part of the ovule (Fig. 1A). The pollen tubes elongated into a depression formed by the female reproductive tissue. In contrast to bryophytes and pteridophytes, which produce abundant sperm in the antheridia, a cycad pollen tube contains only two sperm cells (15). The number of sperm that can be obtained from an ovule is limited to at least a few or at most several dozen depending on the number of pollen tubes. The mature sperm were allowed to exit the pollen tube immersed in a sperm isolation medium (composition shown in Methods; Fig. 1B, C, Movie S1). A 10% sucrose solution described in a previous study (21) was replaced with a medium that reproduced the composition of sexual fluid seeping out of female reproductive tissues just before fertilization. The osmolarity and pH of the medium were adjusted to the actual fluid values (22). Sperm emerging from the pollen tubes gradually activated their flagella and were collected using a pipette. The isolated sperm retained their ability to swim freely in the isolation medium (Fig. 1D, Movie S2). We also observed sperm swimming around the neck cells protruding to the surface of the depression of female tissue filled with the isolation medium (Fig. 1E, Movie S3). The sperm changed direction and repeatedly touched the neck cells as if attracted to the cells. This sperm attraction phenomenon supports previous studies proposing that the sexual fluid from female tissues might contain sperm-attracting molecules secreted from the neck cells (23, 24). These observations indicate that intact motile sperm were isolated. Tissues other than sperm were manually dissected using a razor or tweezers. All collected tissues were processed for sequencing.

**Figure 1.**
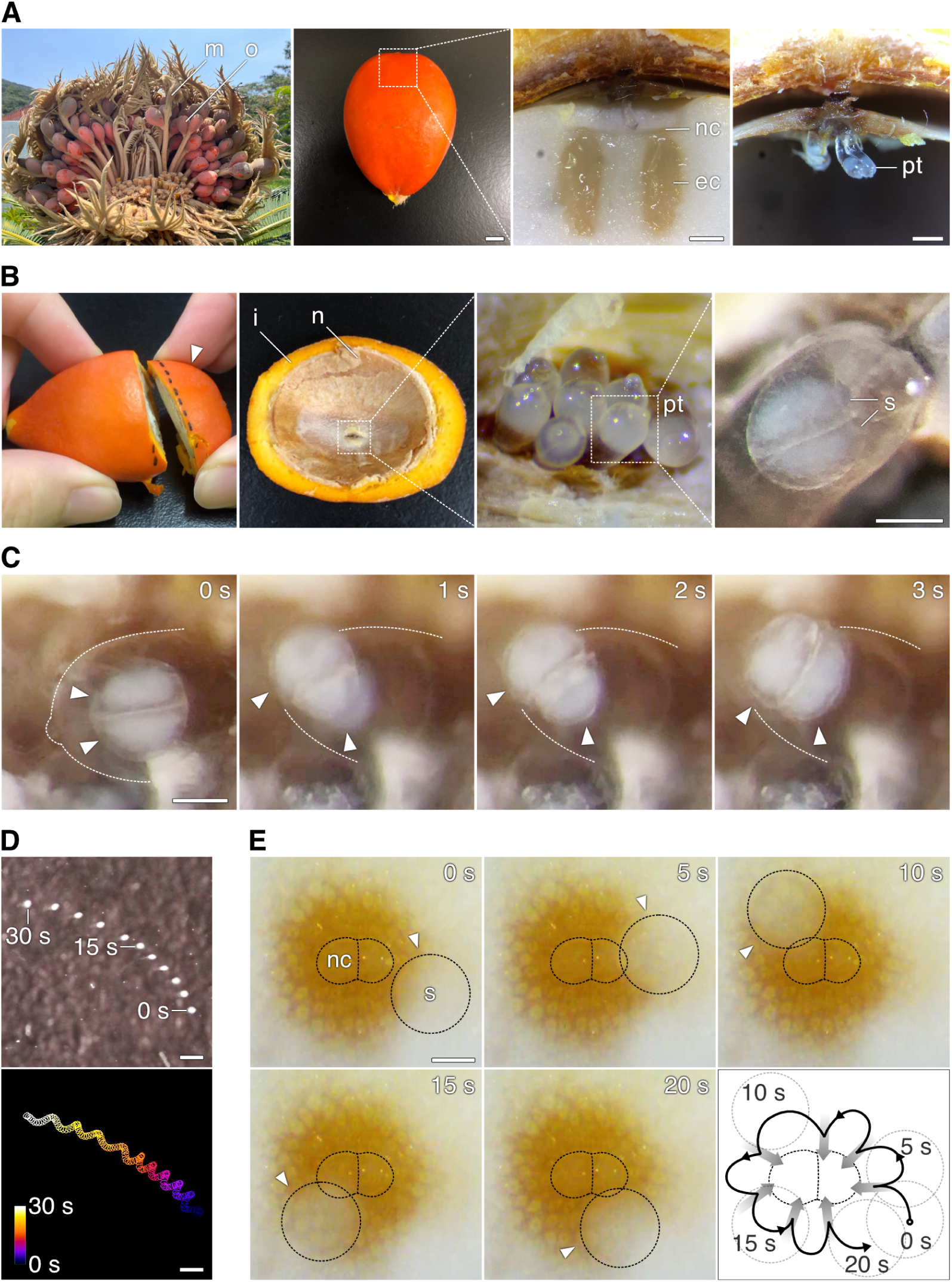
Sperm isolation from ovules and confirmation of sperm viability. (A) Female cone as an aggregate of megasporophylls (m) bearing mature ovules (o). A longitudinal section of the top region of an ovule (white square) shows sexual tissues including neck cells (nc), egg cells (ec), and pollen tubes (pt). (B) Procedure for dissecting ovules to visualize the pollen tubes. Pollen tubes were located at the apical cleavage site of the nucellus (n) next to the integument (i) layer. Two sperm were observed per pollen tube. (C) Selected frames of two sperm exiting a pollen tube (dashed outline). Arrowheads indicate individual sperm cells. (D) Merged images of swimming sperm trajectories every 3 s (upper) and 0.2 s (lower). All frames were binarized to separate sperm and the background; sperm are outlined in each frame (lower). (E) Selected frames at 5-s intervals from a video of sperm swimming around neck cells. Arrowheads indicate sperm positions. Lower right image is a merged image of the sperm trajectory. *Scale bars*, 500 *µ*m (A, D, E) and 100 *µ*m (B, C).

### Control of gene expression in cycad male gametophyte tissues is similar to that in other land plants

To understand the characteristic gene expression dynamics in male cells, cDNA libraries generated from total mRNA from tissues were sequenced on an Illumina platform. Each sample, except one replicate in the megasporophyll, yielded 8–30 million reads (Table 1). For this species lacking publicly available genome data, we performed *de novo* assembly using all sequenced data as inputted in the Trinity program to generate the sequence references. A total of 77,449 protein-coding sequences (CDSs) were obtained, each retaining full length from the start codon to the stop codon (Dataset S1). BUSCO validation for all CDSs was 84–92% complete within universal gene sets of Eukaryotes, Viridiplantae, and Embryophytes, confirming the high quality of the assembled CDSs as reference sequences (Table. S1) (25). To calculate the expression level (transcripts per million, TPM) for each tissue sample, raw sequence data from each tissue was pseudo-aligned to the constructed reference CDSs using the kallisto program (Dataset S2). CDSs with expression > 0 TPM were considered the least detected, and those with expression > 2 TPM were definitively present. Note that “pollen tube with sperm” sample data included expression data from pollen tubes and sperm under differentiation, whereas “sperm” sample data were exclusively from intact sperm.

**Table 1.** RNA sequencing outputs and count data results.

| Tissue Sample | Replicates | Total Sequences | Number of CDSs |  | Tissue Group |
| --- | --- | --- | --- | --- | --- |
|  |  |  | TPM > 0 | TPM > 2 |  |
| Sperm | 1 | 12,045,214 | 38,131 | 9,488 | Male |
|  | 2 | 10,084,787 | 41,145 | 11,690 | Male |
| Pollen tube with Sperm | 1 | 8,316,656 | 62,192 | 24,270 | Male |
|  | 2 | 8,955,365 | 62,153 | 22,736 | Male |
| Neck cell | 1 | 14,756,255 | 65,017 | 30,769 | Female |
| Egg cell with Neck cell | 1 | 26,106,803 | 64,371 | 31,689 | Female |
|  | 2 | 31,900,488 | 66,127 | 31,145 | Female |
| Megasporophyll | 1 | 460,798 | - | - | - |
|  | 2 | 8,752,729 | 42,117 | 21,619 | Flower |
| Leaf | 1 | 14,600,201 | 55,697 | 22,268 | Leaf |
|  | 2 | 26,110,495 | 57,619 | 19,945 | Leaf |
| Integument | 1 | 16,856,119 | 51,909 | 16,481 | - |

In other land plants, transcription is typically suppressed in sperm due to highly condensed nuclear chromatin (14). However, intact cycad sperm exhibited comparable total sequence counts to other tissues, indicating successful mRNA extraction. Additionally, sperm-expressed transcripts covered at least 10% of the total CDSs.

To determine whether the expression patterns of cycad male reproductive cells, including sperm and pollen tubes, differed from those of other land plants, we divided the tissue samples into four groups: male, female, flower, and leaf (Table 1). The expression patterns of the four groups were compared as described in a previous study (26). As in other land plants, male tissues exhibited the lowest CDS expression (Fig. 2A). In contrast, the proportion of specifically expressed CDSs calculated using the specificity measure (SPM) value, which was used as the same threshold in the previous study, was highest in male tissues (Fig. 2B, C). Thus, male-specific CDSs accounted for 7.3% of the total CDSs and 30.7% of CDSs expressed in male tissues (Fig. 2C, Table S2). This result is consistent with other land plants, highlighting that the male-specific CDSs are important genes characterizing their function. In five land plant species, including cycads, we annotated all expressed CDSs using Clusters of Orthologous Genes (COG) categories for functional classification and compared each COG proportion in male tissue-specific CDSs and specific CDSs in pure sperm of *C. revoluta* and *M. polymorpha* or sperm cell of three angiosperms to those in the total CDSs (Fig. 2D). The proportion of genes involved in information storage, such as transcription, translation, and associated post-translational functions decreased in all species, indicating similar transcriptional suppression to that in other land plants. Although the cycad sperm nucleus is not condensed during development, transcriptional repression likely occurs similarly to other land plant species (15, 19). However, genes related to the cytoskeleton were consistently enriched, probably involving the formation of the flagella in motile sperm and growth of the tip and inner cell migration of the pollen tubes (27, 28). Additionally, signal transduction, which may be related to male and female interaction during fertilization, had the highest proportion in male tissues and sperm from four seed plant species, including the cycad, and the proportion was higher than in all other tissues. This finding suggests that even in cycads that use motile sperm, some transitions may have occurred in the reception of female signals and signal transduction within male reproductive cells. The proportion of cell wall/membrane/envelope biogenesis increased in only three angiosperm species, suggesting the emergence of novel factors for sperm cell traits or angiosperm-like pollen tube traits. These similarities and differences in the trends of these COG categories reflected the molecular mechanisms controlling cycad sperm and pollen tube functions.

**Figure 2.**
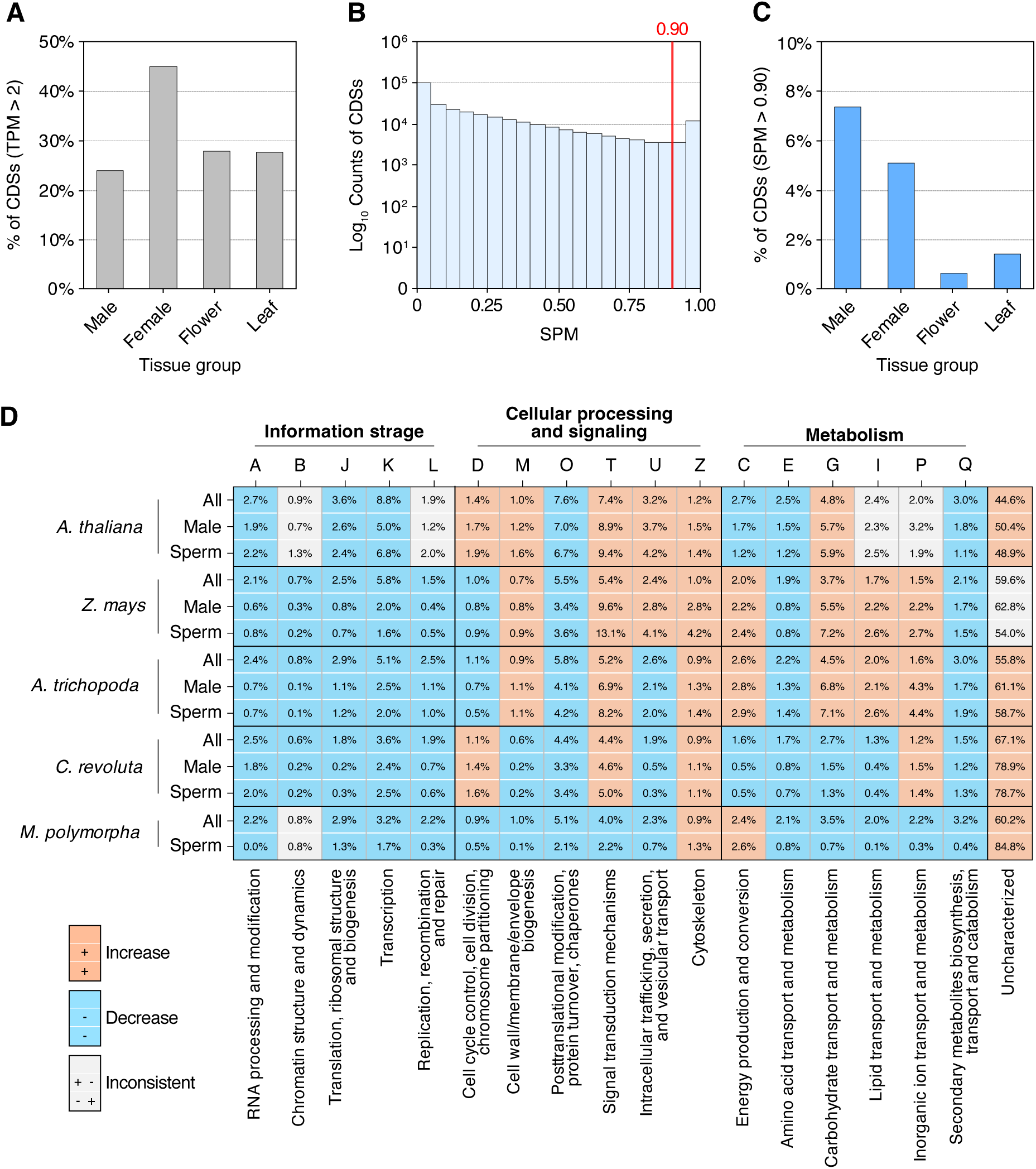
(A) Percentage of protein-coding sequences (CDSs) (y-axis) with expression levels > 2 transcripts per million (TPM) in each tissue group (x-axis) relative to the number of total CDSs. (B) Histogram showing the distribution of specificity measure (SPM) values (x-axis) calculated in each tissue for all CDSs. The number of CDSs in each bin was converted to a logarithmic scale (y-axis). Specifically expressed CDSs with the top 5% SPM values are shown above the red line (SPM = 0.90). (C) Percentage of specifically expressed CDSs (y-axis) (SPM > 2) in each tissue group (x-axis) among all CDSs. (D) Percentages of CDSs annotated into each Clusters of Orthologous Genes (COG) category (horizontal row) for five land plant species, including *Cycas revoluta* (vertical column). Descriptions for each COG category are provided in the lower row. Values for all expressed genes (“All”, upper row), male tissue-specific expressed genes (“Male”, middle row), and sperm-expressed genes (“Sperm”, lower row) are shown for each species. For each COG category in a certain species, if “Male” and “Sperm” values exceeded the “All” value, the three corresponding boxes are red; otherwise, they are blue.

### Expression patterns of histone variants indicate histone substitution in cycad sperm

A representative mechanism regulating transcriptional repression in male reproductive cells involves replacing somatic histone variants with SNBPs during spermatogenesis, leading to chromatin condensation (29). This mechanism is common in animals and plants and shows the lineage-specific evolution of the SNBP types (4, 30). Bryophyte sperm mainly use a protamine-like protein type (PL type), similar to that of many metazoan sperm (31). Pteridophyte sperm use both histone types (H type) and protamine types (P type), whereas angiosperms completely lack the P and PL types, but use the H type (5, 32). Studies of SNBPs in cycads remain limited, and no protamine types typical of flagellated sperm have been identified. To investigate SNBP status in cycads, we identified histone-variant (H1, H2A, H2B, and H3) homologs, previously reported for their sperm specificity in other plants, in a BLAST search. Variants specifically expressed in the sperm of some angiosperm species have been detected for histones H2A, H2B, and H3, which are associated with chromatin remodeling (30, 33, 34). In *Marchantia polymorpha*, protamine is produced by post-translational modification of sperm-specific H1 histones, i.e., PL-type SNBPs (31). Of the 77,449 CDSs, the expression patterns of the candidate homologs were visualized in heatmaps (Fig. 3, Tables S3 and S4), and phylogenetic trees were constructed for their classification (Figs. S1–S4, Tables S5–S8).

**Figure 3.**
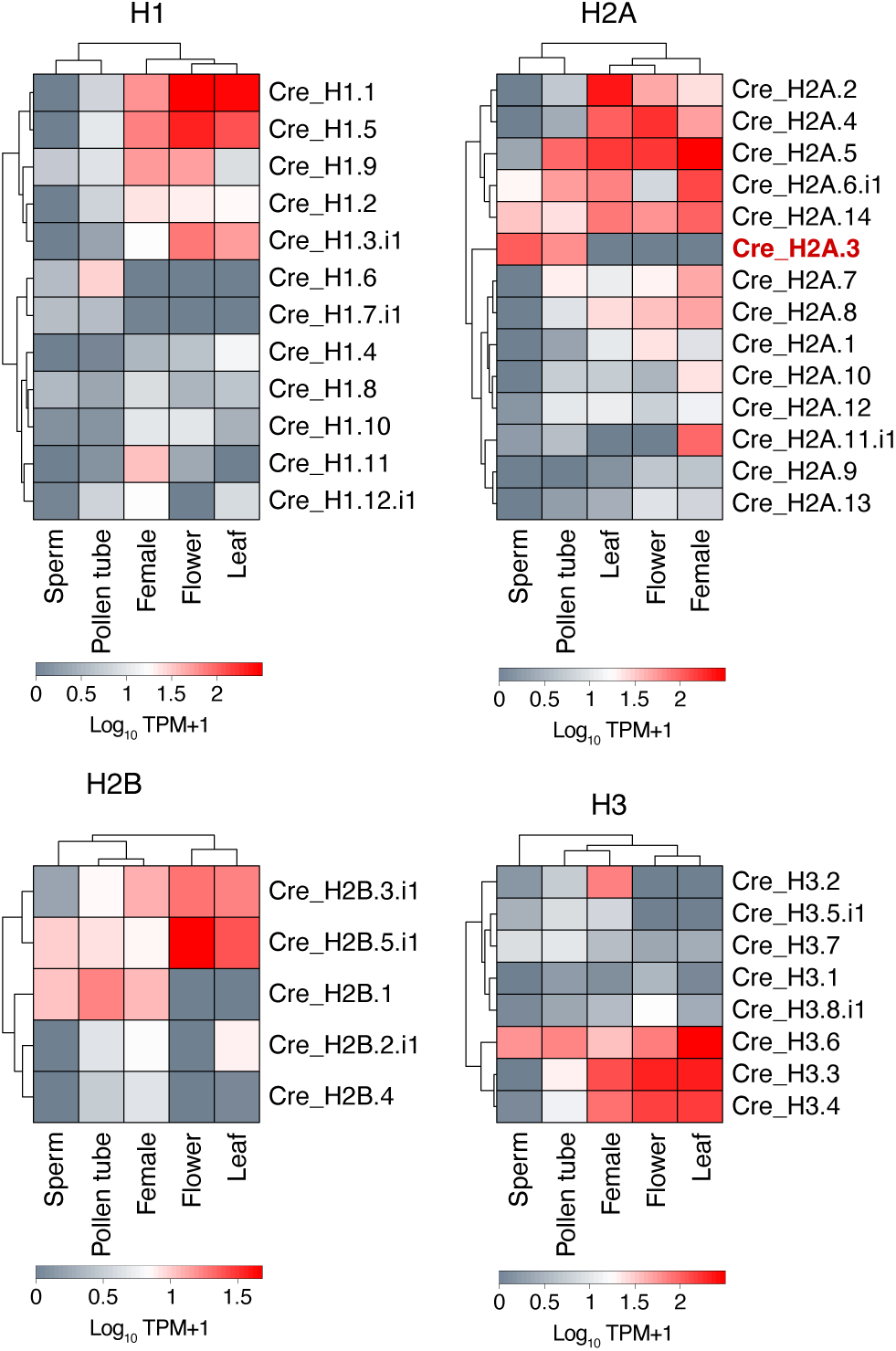
Heatmaps showing expression patterns of the candidate *C. revoluta* CDSs for the histone variants H1, H2A, H2B, and H3. Vertical axis lists the identified isoforms corresponding to each histone variant; horizontal axis represents each tissue or tissue group. Expression levels are log-scaled. Isoforms marked in red indicate exclusive expression in sperm and pollen tubes.

Several constitutively expressed histones within each H1, H2A, H2B, and H3 histone variant were significantly downregulated in sperm and pollen tubes, suggesting the loss of somatic histones in male reproductive cells. In contrast, the most sperm-specific histone was “Cre_H2A.3”, which was classified as H2A.W in the H2A phylogenetic tree, potentially defining the heterochromatin region (35). Within H2A, downregulated variants in sperm such as “Cre_H2A.7”, “Cre_H2A.1”, and “Cre_H2A.12” were classified as H2A.Z, and associated with protecting the euchromatin (36). H2A variants displayed an expression profile that promotes chromatin condensation in sperm nuclei. Although “Cre_H1.6” was also male-specific, its expression was higher in pollen tubes than in sperm, and its sequence did not contain abundant arginine, a characteristic of protamines. This result indicates that cycad sperm has no P-type variants and relies on an H-type SNBP for its transcriptional repression mechanism, as observed in some pteridophyte and angiosperm sperm cells. Despite the large cell and nuclear sizes of cycad sperm, our results suggest that transcriptional repression in cycad sperm is controlled by replacing somatic histones with SNBP as well as male gametes in other plants.

### Conservation of fertilization-related cell membrane-localized genes in cycad sperm

Among the CDSs classified in the COG signal transduction category, we focused on membrane-localized genes involved in male–female interactions. The proportion of genes predicted to be localized to the plasma membrane among male reproductive cell-specific CDSs was lower than the overall proportion, and no difference was observed between lineages (Fig. 4A). Functional classification of factors predicted to be membrane-localized revealed differences in their composition among lineages (Fig. 4B, Table S9). Cycads and *M. polymorpha* had only one or two RLKs, whereas the three angiosperm species had more RLKs and the number of leucine-rich repeat-RLKs (LRR-RLKs) not found in cycads or bryophytes was the highest in all three species, suggesting the acquisition of novel RLKs in angiosperms (37). This finding suggests a change in the mechanism by which motile and non-motile sperm receive female signals. Focusing on non-receptor genes, the potassium ion channel (AKT) and CDC50 were common to the three angiosperm species and cycads. CDC50, also known as aminophospholipid ATPase (ALA-interacting subunits, ALIS), forms P4-ATPase complexes with ALA, which functions as a flippase, catalyzing the flipping of cell membrane phospholipids (38). These factors indicate a common mechanism for regulating ion concentrations and cell membrane conditions in male gametes between angiosperms and cycads (39–41). A glutamate receptor-like channel (GLR) and CAPE, which have fertilization-related functions in bryophytes and pteridophytes, were also detected in the cycad. GLR has functions in sperm chemotaxis in the bryophyte *Physcomitrella patens* and in pollen tube development, suggesting that the cycad GLR may have similar functions (11, 42). CAPE is distributed only in streptophytes with motile sperm and is involved in the control of sperm motility through cAMP signaling, providing solid evidence of motile sperm characteristics in cycads (43, 44).

**Figure 4.**
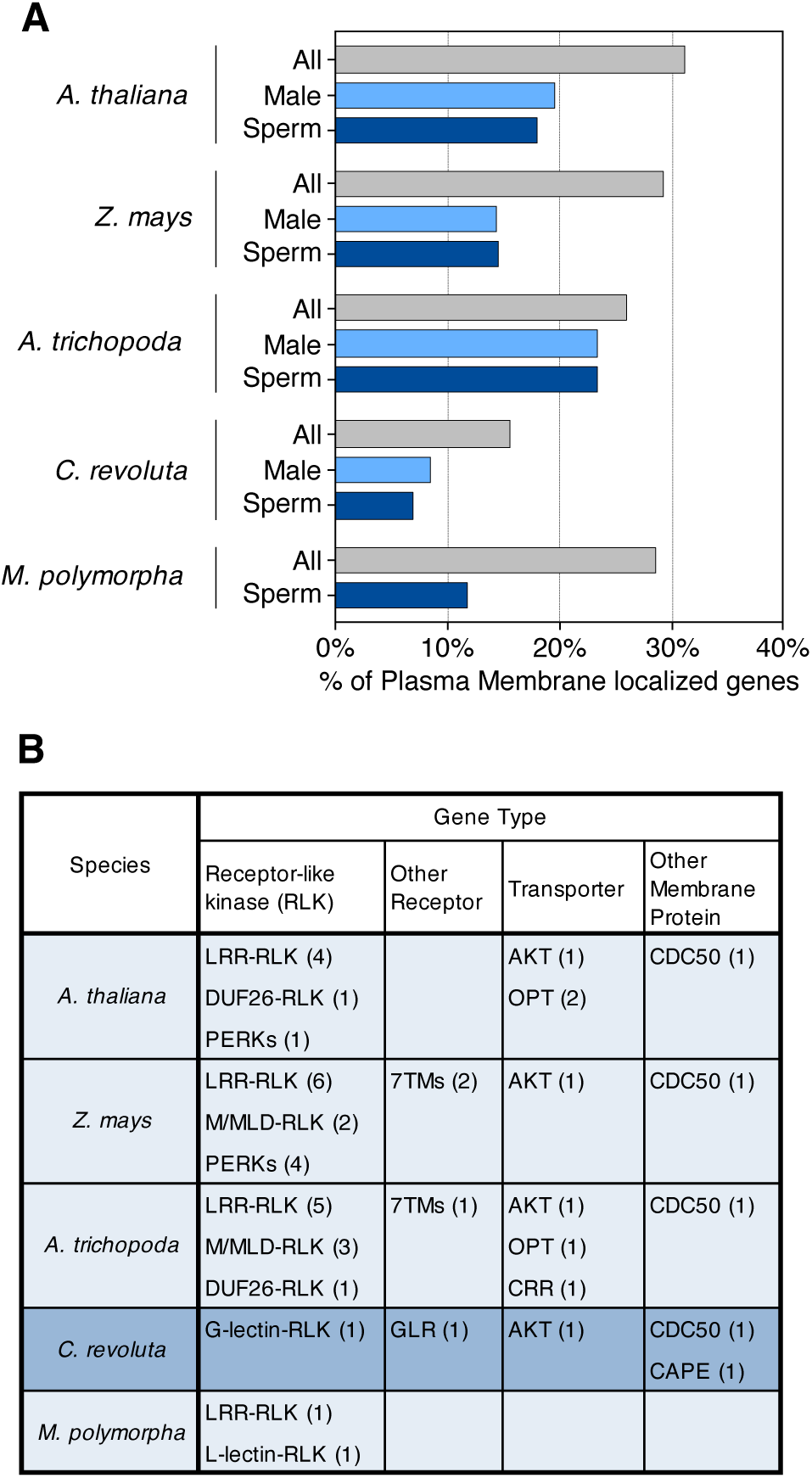
Gene type transition of sperm-specific genes associated with signal transduction. (A) Percentage of genes predicted to localize to the plasma membrane within genes associated with signal transduction (COG = T) for each species. (B) Gene type classification of sperm-specific genes predicted to localize to the plasma membrane. Each gene abbreviation includes genes in the same families; the number of genes is indicated in parentheses.

Next, we identified putative plasma membrane-localized factors by predicting signal peptides and transmembrane domains, even for factors not associated with signal transduction according to COG annotation (Table S10). Factors involved in gamete fusion, such as Hapless 2/Generative Cell Specific 1 (HAP2/GCS1) and Gamete Expressed 2 (GEX2), indicated that the gamete fusion mechanism is conserved in cycads (8, 45). Transporters were found most frequently, whereas only two RLKs were found. Although the functions of the homologous genes of these RLKs have not been reported in angiosperm fertilization, they may receive the female signal during fertilization in cycads. These results suggest that cycad sperm have partially acquired sperm cell-like molecules functioning in angiosperms while retaining molecules required for motile sperm. It remains necessary to verify which factors function in the interaction between cycad sperm and female signals by functionally analyzing these candidate molecules.

### Angiosperm pollen tube-like functions are incomplete in cycad pollen tubes

Many of the putative plasma membrane-localized factors expressed in sperm were also expressed in pollen tube samples containing sperm; however, there were no homologs of already-known RLKs that function in receiving female signals during pollen tube attraction in angiosperms. To verify the expression dynamics of known RLKs specifically expressed in the pollen tubes of angiosperms, we investigated the expression and phylogenetic relationships of these genes, including malectin and malectin-like domains (M/MLD) in cycads. FERONIA (FER) is a representative gene in female angiosperm reproductive tissues; among genes with the same domain structure (MLD-RLK) with FER, ANXUR 1/2 (ANX1/2) and Buddha’s Paper Seal 1/2 (BUPS1/2) are expressed specifically in pollen tubes and are involved in pollen tube integrity (46, 47). In contrast, *M. polymorpha* has no pollen tube and has a single copy of FER expressed ubiquitously in various tissues (48). Five candidates containing M/MLD were detected within male reproductive cell-specific CDSs of the cycad, none of which exhibited male-specific expression (Fig. 4A, Table S11). Phylogenetic analysis showed that only one of five candidates was classified into the MLD-RLK group: *M. polymorpha* (Fig. 4B, Table S12), suggesting that M/MLD-containing genes associated with pollen tube guidance evolved in the angiosperm lineage and have not yet diversified in cycads.

COG classification identified common factors such as exostosins and mechanosensitive ion channels as pollen tube formation factors in the biogenesis of cell walls, membranes, and envelopes (Table S13). However, some factors were common among angiosperm pollen tubes, but not present in cycads. Typical examples included callose synthase and UDP-D-glucuronate 4- epimerase, which are involved in organization of the cell wall components callose and pectin (49, 50). Additionally, galactolipid monogalactosyldiacylglycerol, which functions during pollen tube elongation, is detected only in angiosperms (51). These factors have been described as important for the rapid growths of pollen tube tips. The finding that they were absent in the cycad is consistent with previous observations that cycad pollen tubes have not acquired the mechanism for tip growth (17, 49, 52). It appears that cycad pollen tubes lack key functions such as guidance to egg cells via female signals and rapid tip growth, which enable pollen tubes to achieve siphonogamy, as observed in angiosperms.

## Discussion

In this study, we successfully obtained sufficient quantities of ovules during fertilization by continuously observing fertilization in *C. revoluta*. We also developed a method to collect cycad sperm by creating a sperm isolation medium that closely mimics the actual fertilization fluid by adjusting osmolarity and pH (Fig. 1). This medium allowed us to isolate sperm despite the limited number of cells that can be obtained from a single ovule. This collection method enabled the first transcriptome analysis of mature cycad sperm. We conducted RNA sequencing of several cycad tissue samples, including reproductive and non-reproductive tissues (Table 1). The reference sequences of transcripts (77,449 CDSs) constructed through *de novo* assembly using the sequenced data exhibited sufficient quality, as indicated by the BUSCO value (Table S1). Mapping to the reference sequence enabled the quantification of gene expression levels across tissues, allowing their comparison (Table 1). Total reads in mature sperm showing full pollen tube emergence and swimming were found to be comparable with those of other tissues; their expression included 10% of all reference CDSs. Cycads have uniquely large-volume, multi-flagellate sperm and large pollen tubes that are morphologically very different from those of other land plants. We focused on the gene expression dynamics of cycad sperm and pollen tubes, and fertilization-related factors contributing to interactions with female reproductive tissues.

### Relationship between the control of gene expression and sperm nuclear basic proteins in cycad sperm

In contrast to the few genes expressed in cycad male reproductive tissues, the proportion of specifically expressed genes was remarkably high, whereas genes related to transcription and translation were strongly suppressed (Fig. 2). This tendency is consistent with other land plants, suggesting an underlying gene expression control program conserved across the land plant lineage (26). Therefore, we focused on SNBPs as a representative factor controlling gene expression in male gametes common to plants and animals. The putative SNBP “Cre_H2A.3” (H-type) was found to be a histone specifically expressed in cycad sperm (Figs. 3 and S1–S4). SNBP types have transitioned several times in animals, and recent studies have reported a transition of the H type in plant lineages (4, 5). Our finding is a key element in bridging the gap between basal land plants and angiosperms, leading us to hypothesize that the transition from the PL type to the H type has occurred in a land plant lineage, specifically in pteridophytes. Despite the morphological uniqueness of cycad male gametophyte tissues, such as large cells and nuclei, these results indicate a commonality in the transcriptional regulatory system.

However, unlike the minimal transcription observed in other land plant sperm nuclei caused by high nuclear condensation, a particular level of gene expression was detected in cycad sperm (Table 1) (1). We hypothesized that this result reflected reduced nuclear condensation, allowing residual transcriptional activity, and questioned whether transcription is minimal, as in other plant sperm but with transcripts pre-accumulated during spermatogenesis. Few detailed studies have examined the degree of nuclear condensation in cycads, partly because of a specific step in which generative cells in mature pollen are transformed into exceptionally large size sperm (10 times that of generative cells) over a long period (about 3 months in *C. revoluta*) (53). This significant size difference complicates direct estimates of the level of cycad sperm nuclear condensation through examining generative cells. Such comparisons are simpler in angiosperms, where generative cells and sperm cells are similar in size and have similar levels of nuclear condensation (54). Future work will require visualizing the transcriptionally active regions in sperm and pollen tube cells, analyzing the expression and localization of sperm-specific histone variants, and epigenetic profiling of the variants. Additionally, the loose nuclear chromatin condensation in cycad and *Ginkgo* sperm may represent an adaptive feature of motile sperm in seed plants. It is necessary to verify how this sperm morphology and the expression dynamics of seed plants are adaptive in the environment within the ovule where the sperm swim and in interactions with female reproductive tissues. It will also be necessary to clarify the differences between cycad sperm and the sperm of basal land plants.

### Fertilization-related genes provide insight into the evolutionary trajectory of sperm motility loss and pollen tube acquisition

A search for candidate membrane-localized factors in sperm and pollen tubes revealed that cycad sperm possess membrane factors common to motile sperm from basal land plants and non-motile sperm cells from angiosperms. In contrast, pollen tubes lack female receptor factors necessary to trigger the attraction of the pollen tube and wall components, which are important in tip growth (Figs. 4 and 5, Tables S9–S13). The expression profile of cycad pollen tubes did not support a specialized pollen tube function, as observed in angiosperms; it is likely that the cycad pollen tube has only a haustorial function, as suggested by previous anatomical analyses (17). These results demonstrate the molecular intermediate state of fertilization in cycads using motile sperm and pollen tubes. It is possible that such an intermediate state occurred when a fertilization system using non-motile sperm and pollen tubes was acquired in the angiosperm lineage. These results further provide important molecular information for exploring the evolutionary process leading to siphonogamy. In contrast, extant gymnosperms are monophyletic (55–57). The fertilization expression profile of cycads may be a direct intermediate state in the process of siphonogamy establishment in gymnosperm lineages. Future studies are required to compare these expression profiles with those of the reproductive cells of conifers and gnetales.

**Figure 5.**
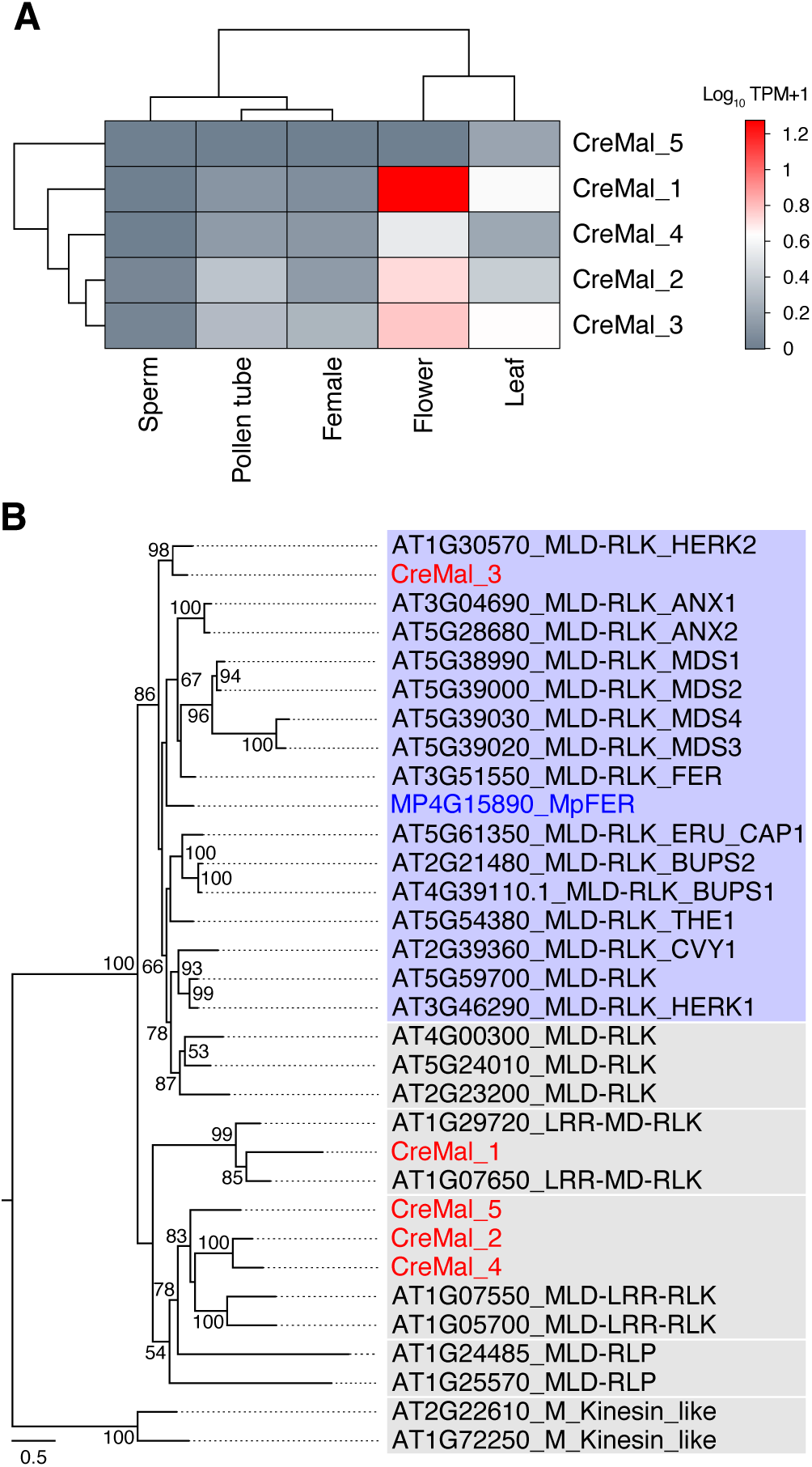
Identification and expression patterns of CDSs containing malectin or malectin-like domains (MD/MLD). (A) Expression patterns of five *C. revoluta* MD/MLD containing CDSs. Vertical axis lists the five CDSs; horizontal axis represents each tissue or tissue group. (B) Maximum-likelihood phylogenetic tree constructed from the protein sequences of all genes or CDSs containing M/MLD across *Arabidopsis thaliana*, *C. revoluta*, and *Marchantia polymorpha* (1,000 bootstrap replicates). Red genes indicate *C. revoluta*, blue genes indicate *M. polymorpha*; all others are from *A. thaliana*.

Because gamete fusion factors are conserved, it is possible that evolutionary variation arose among lineages in the molecular interactions between males and females before gamete fusion. Changes in the attributes of female signals received by sperm and pollen tubes to guide them to the egg cell are accompanied by changes in the morphology and plasma membrane-localized molecules of male reproductive cells in the land plant lineage. For example, bryophyte and pteridophyte sperm receive inorganic ions and hydrocarbons as chemotactic molecules, whereas RLKs in angiosperm pollen tubes receive peptide molecules (13, 58, 59). Therefore, it is important to determine which cell membrane-localized proteins of cycad sperm are functional and which female signals are received to guide the sperm to the egg cell, to infer the evolution of the molecular mechanisms of male and female interactions preceding gamete fusion. Future research will elucidate the detailed evolutionary state of cycad fertilization by analyzing the interactions between signal molecules derived from female reproductive tissues and the cell membrane-localized candidate factors in cycad sperm identified in this study.

## Materials and Methods

### Plant materials

We selected three *Cycas revoluta* Thunb. habitats in Japan where male and female trees coexisted: Iriomote Island in Okinawa, Kibana campus of the University of Miyazaki in Miyazaki, and the Koishikawa Botanical Gardens of The University of Tokyo in Tokyo. Female cones were artificially pollinated using mature pollen collected from male cones. Three months after pollination, progress to fertilization was monitored daily in each habitat for about 2 weeks, and mature ovules were sampled almost at the point of fertilization.

### Sperm isolation

The top 1 cm of the ovule was cut with pruning scissors to expose the germline cells. The orange outer segment included the integument and nucellus with pollen tubes. The white inner segment was the female gametophyte, including the neck and egg cells. To isolate the sperm, about 100 *µ*L of sperm isolation medium (5 mM Tris-HCl, 2 mM KOH, pH 8.4, 0.83 M D(+)-glucose, 0.11 M D(-)-fructose) was added to the bottom of the nucellus, where pollen tubes were protruding (22). The intact mature sperm swam out from the pollen tube immersed in the medium and were caught with a pipette (21).

### Stereomicroscopic observations

All microscopic images were obtained by an SZX7 stereomicroscope (0.1 numerical aperture, 1× objective) (Olympus, Tokyo, Japan) and captured using a 12MP Wide iPhone 8 camera attached to an eyepiece lens via an i-NTER LENS [MR-6i] (Micronet, Tokyo, Japan) (Fig. 1A–C). Movies were recorded at 30 fps with a 4-K video recording tool using the same camera (Fig. 1D, E). Merged images shown in Fig. 1D were processed with the Z-projection program in Stacks and the Temporal-Color Code program in Hyperstacks (ImageJ/Fiji, National Institutes of Health, Bethesda, MD, USA).

### RNA isolation and sequencing

Tissues for RNA sequencing were collected from *C. revoluta* trees grown outdoors at the Koishikawa Botanical Gardens. Four types of tissues containing male and female germline cells and three types of germline-free tissues were included. Two types of male samples were collected: ten isolated sperm and ten pollen tubes containing sperm. Two types of female samples were dissected with a razor blade from female gametophyte tissue: three tissue pieces containing neck cells, and three pieces containing an egg cell and neck cells. Three types of germline-free tissues were collected from female *C. revoluta* trees: 5 mg each of megasporophyll, mature leaves, and integument. Each tissue sample was collected into a 1.5-mL DNA LoBind Tube (0030108051, Eppendorf, Mannheim, Germany), and total mRNA was extracted with a Dynabeads mRNA DIRECT micro kit (no. 61021, ThermoFisher Scientific, Waltham, MA, USA) according to the manufacturer’s protocol. The cDNA libraries for each sample were prepared with the TruSeq RNA Sample Prep Kit v2 (nos. 15027387, 15025063, and 15027084, Illumina, San Diego, CA, USA), following the low-sample protocol of the manufacturer. The replicate numbers per sample are listed in Table 1. Twelve libraries with individual adapters were pooled and sequenced into four lanes on the Illumina NextSeq 500 platform to generate paired-end reads.

### CDS dataset construction and gene expression profiles

FASTQ files for the sequenced data were generated with Illumina bcl2fastq software. Bases with a quality value < 20 were replaced with N in each read, and reads containing ≥ 40 consecutive non-N bases were retained. To acquire all of the transcript sequences, *de novo* assembly was performed using Trinity v2.11.0, with a minimum length of 300 bases and inputting all trimmed FASTQ reads (60). The reference CDSs were detected using TransDecoder v5.5.0 with default settings from the Trinity output, and the sequences annotated “complete” were extracted (Dataset S1). To check the quality of the 77,449 CDSs, the completeness of the core gene set was quantified using BUSCO v5.5.0 (Table S1).

The expression levels of all CDSs were estimated using kallisto v0.46.2 with default settings to pseudo-align the FASTQ reads of each sample, except for the low-quality megasporophyll replicate1, to an index created from the total CDSs (Dataset S2) (61). To compare the tissue-specific gene expression profiles in *C. revoluta* with other plant lineages, the sequenced samples were divided into four tissue groups as described in a previous study (26): Male (“sperm” samples and “pollen tubes with sperm” samples), female (“neck cell” samples and “egg cells with neck cells” samples), flower (megasporophyll, only replicate 2), and leaf (leaf). CDSs with an average of > 2 TPM were distinctly expressed (Table 1, Fig. 2A). To find tissue-specific expressed CDSs, SPM values were calculated as described previously. CDSs with SPM > 0.90, where 95% of all expressed CDSs were found, were defined as differentially expressed CDSs (Fig. 2B, C, Table S2). The eggNOG-mapper tool was used to annotate the COG categories for each CDS to compare the proportions of COG annotations for all genes expressed in the plant and for genes specifically expressed in male tissues plus sperm or sperm cells (expression > 2 TPM, particularly in sperm or sperm cells) (Fig. 2D). CDSs and the *C. revoluta* expression data were obtained in the present study, and data for other plants were obtained from a previous study (26).

### Homology inference and phylogenetic analysis of histone variants

BLAST+ was used to detect the CDSs of the histone variant candidates. The total CDSs of *C. revoluta* were used as the database, whereas representative histone variant sequences from *Arabidopsis thaliana* (Ath), *Marchantia polymorpha* (Mpo), and *Volvox carteri* (Vca) were used as query sequences, including Ath_H1.1 and Mpo_H1.1 for H1, Ath_H2A.1 and Mpo_H2A for H2A, Ath_H2B.1, Mpo_H2B.1, and Vca_H2B.13 for H2B, and Ath_H3.1 and Vca_H3-like_1 for H3. The candidate CDSs were further verified using the National Center for Biotechnology Information (NCBI) Web CD-Search Tool. Only CDSs containing conserved domains annotated as histones and with e ≤ 1^−6^ were retained (Table S3). Heatmaps for all *C. revoluta* histone variant candidates were analyzed using the R package *pheatmap* v1.0.12 (Fig. 3). The raw TPM values are listed in Table S4. The sequence and expression patterns of the i1 isoform for histones with multiple isoforms were used to generate phylogenetic trees and heatmaps. Phylogenetic trees were constructed to analyze the relationships of these histone-predicted CDSs in comparison with those from other species (Figs. S1–S4). Sequence alignments were generated using MAFFT v7 and trimmed with trimAl v1.4.1 to retain the most reliable positions. Maximum likelihood trees were created with RAxML-NG v1.2.1, employing 1,000 bootstrap replicates based on the model determined by ModelTest-NG v0.1.7.

Histone protein sequences used for the blast search and the phylogenetic trees were primarily obtained from previous studies, as indicated in the Supplementary Information. Additional sequences were retrieved from Phytozome and MarpolBase (Tables S5–S8).

### Prediction of CDSs localized to sperm or the sperm cell plasma membrane

To predict the localization of genes annotated to the signal transduction COG category, we used DeepLoc v2.0, SignalP v6.0, and TMHMM v2.0. For each of the five species, genes whose DeepLoc annotation predicted locations and membrane types were cell membrane and transmembrane and genes predicted to be signal peptides and to have one or more transmembrane regions were extracted as candidate genes localized to the plasma membrane (Fig. 4A).

To search for plasma membrane-localized factors in genes predicted not to be related to signal transduction (COG categories other than “T”) in *C. revoluta*, we used the same method to extract candidate factors from 5,679 CDSs (> 2 TPM, SPM > 0.90) and 841 CDSs (0 TPM < expression ≤ 2 TPM, SPM = 1) specifically expressed in male tissues. The gene types were predicted with a BLAST search (Fig. 4B, Tables S9 and S10).

### Screening and phylogenetic analysis of CDSs with malectin or malectin-like domains

Among all CDSs, 1,180 had at least one transmembrane region predicted by TMHMM v2.0 and a signal sequence predicted by Signal-P v6.0. Of these, five CDSs containing malectin M/MLD were screened using the NCBI Web CD-Search Tool, excluding duplicates (Table S11). The phylogenetic tree and heatmap of the expression pattern were constructed as described for histones (Fig. 5). The accession numbers of these genes are listed in Table S12.

## Data availability

Data deposition: The sequences and expression data of *de novo* assembled CDSs are included as Dataset S1 and S2. Raw RNA sequencing reads are available at NCBI Sequence Read Archive (accession no. PRJNA1240061).

## Acknowledgments

The authors thank Dr. Tokushiro Takaso (Iriomote Island on Okinawa) for providing *Cycas revoluta* Thunb. plant material and for advice. We thank the Koishikawa Botanical Garden (The University of Tokyo) for providing the cycad plant materials. We also thank Ayami Furuta for technical assistance. This study was supported by Japan Society for the Promotion of Science (JSPS) Grants-in-Aid (no. 23KJ0741 to Research Fellow Y.T.; nos. 22H05172 and 22H05178 for Transformative Research Areas [A] to S.O.; and no. 22H04980 for Scientific Research [S] to T.H.) and by a Key-Molecule-Network in Plant Reproduction (KEPLR) International Leading Research Grant-in-Aid (no. 22K21352 to T.H.).

The English in this document has been checked by at least two professional editors, both native speakers of English. For a certificate, please see: http://www.textcheck.com/certificate/ZKwaaE

## Author Contributions

Y.T., S.O., and T.H. designed research, Y.T. performed research, Y.T. and T.S. collected data, Y.T. analyzed data, Y.T. wrote the original draft, S.O., T.S., and T.H. reviewed and edited original draft, and T.H. supervised.

## Competing Interest Statement

Disclose any competing interests here.

## Classification

Biological Sciences, Plant Biology

## Notes

https://www.ncbi.nlm.nih.gov/bioproject/PRJNA1240061/

### Competing Interest Statement

The authors have declared no competing interest.

## References

1. D. Hackenberg, D. Twell, The evolution and patterning of male gametophyte development. Curr. Top. Dev. Biol. 131, 257–298 (2019).

2. W. E. Friedman, The evolutionary history of the seed plant male gametophyte. Trends Ecol. Evol. 8, 15–21 (1993).

3. H. Nishida, K. B. Pigg, K. Kudo, J. F. Rigby, Zooidogamy in the Late Permian genus Glossopteris. J. Plant Res. 117, 323–328 (2004).

4. J. M. Eirín-López, J. Ausió, Origin and evolution of chromosomal sperm proteins. Bioessays 31, 1062–1070 (2009).

5. J. Ausió, et al., The sperm nuclear basic proteins of the sword fern (Polystichum munitum). Biochem. Cell Biol. 102, 285–290 (2024).

6. A. Higo, et al., Transcription factor DUO1 generated by neo-functionalization is associated with evolution of sperm differentiation in plants. Nat. Commun. 9, 5283 (2018).

7. S. Yamaoka, et al., Generative cell specification requires transcription factors evolutionarily conserved in land plants. Curr. Biol. 28, 479–486.e5 (2018).

8. T. Mori, H. Kuroiwa, T. Higashiyama, T. Kuroiwa, GENERATIVE CELL SPECIFIC 1 is essential for angiosperm fertilization. Nat. Cell Biol. 8, 64–71 (2006).

9. T. Mori, T. Igawa, G. Tamiya, S.-Y. Miyagishima, F. Berger, Gamete attachment requires GEX2 for successful fertilization in Arabidopsis. Curr. Biol. 24, 170–175 (2014).

10. A. Higo, et al., Transcriptional Framework of Male Gametogenesis in the Liverwort Marchantia polymorpha L. Plant Cell Physiol. 57, 325–338 (2016).

11. C. Ortiz-Ramírez, et al., GLUTAMATE RECEPTOR-LIKE channels are essential for chemotaxis and reproduction in mosses. Nature 549, 91–95 (2017).

12. N. Luo, et al., Exocytosis-coordinated mechanisms for tip growth underlie pollen tube growth guidance. Nat. Commun. 8, 1687 (2017).

13. Y. Mizuta, T. Higashiyama, Chemical signaling for pollen tube guidance at a glance. J. Cell Sci. 131 (2018).

14. D. Southworth, M. Cresti, Comparison of flagellated and nonflagellated sperm in plants. Am. J. Bot. 84, 1301 (1997).

15. K. S. Renzaglia, D. J. Garbary, Motile Gametes of Land Plants: Diversity, Development, and Evolution. CRC Crit. Rev. Plant Sci. 20, 107–213 (2001).

16. L. Alvarez, The tailored sperm cell. J. Plant Res. 130, 455–464 (2017).

17. B. M. Johri, Haustorial Role of Pollen Tubes. Ann. Bot. 70, 471–475 (1992).

18. The Effects of Fertilization on the Growth andPhysiological Characteristics of Ginkgo biloba L.

19. K. J. Norstog, T. J. Nicholls, The biology of the cycads (Cornell Univ. Press, 1997).

20. E. Offer, S. Moschin, S. Nigris, B. Baldan, Reproductive Mechanisms in Ginkgo and Cycas: Sisters but not Twins. CRC Crit. Rev. Plant Sci. 42, 283–299 (2023).

21. T. Hori, S.-I. Miyamura, “Contribution to the Knowledge of Fertilization of Gymnosperms with Flagellated Sperm Cells: Ginkgo biloba and Cycas revoluta” in Ginkgo Biloba A Global Treasure: From Biology to Medicine, T. Hori, et al., Eds. (Springer Japan, 1997), pp. 67–84.

22. P. von Aderkas, et al., Composition of Sexual Fluids in Cycas revoluta Ovules During Pollination and Fertilization. Bot. Rev. 88, 453–484 (2022).

23. D. Wang, et al., Structure and function of the neck cell during fertilization in Ginkgo biloba L. Trees 28, 995–1005 (2014).

24. T. Takaso, Y. Kimoto, J. N. Owens, M. Kono, T. Mimura, Secretions from the female gametophyte and their role in spermatozoid induction in Cycas revoluta. Plant Reprod. 26, 17–23 (2013).

25. V. Raghavan, L. Kraft, F. Mesny, L. Rigerte, A simple guide to de novo transcriptome assembly and annotation. Brief. Bioinform. 23 (2022).

26. I. Julca, et al., Comparative transcriptomic analysis reveals conserved programmes underpinning organogenesis and reproduction in land plants. Nat Plants 7, 1143–1159 (2021).

27. J. Lucas, M. Geisler, Sequential loss of dynein sequences precedes complete loss in land plants. Plant Physiol. 189, 1237–1240 (2022).

28. Y. Fu, The cytoskeleton in the pollen tube. Curr. Opin. Plant Biol. 28, 111–119 (2015).

29. J. Ausió, M. L. Van Veghel, R. Gomez, D. Barreda, The sperm nuclear basic proteins (SNBPs) of the sponge Neofibularia nolitangere: implications for the molecular evolution of SNBPs. J. Mol. Evol. 45, 91–96 (1997).

30. T. Buttress, et al., Histone H2B.8 compacts flowering plant sperm through chromatin phase separation. Nature 611, 614–622 (2022).

31. R. A. D’Ippolito, et al., Protamines from liverwort are produced by post-translational cleavage and C-terminal di-aminopropanelation of several male germ-specific H1 histones. J. Biol. Chem. 294, 16364–16373 (2019).

32. W. F. Reynolds, S. L. Wolfe, Protamines in plant sperm. Exp. Cell Res. 152, 443–448 (1984).

33. K. Ueda, et al., Male gametic cell-specific histone gH2A gene of Lilium longiflorum: genomic structure and promoter activity in the generative cell. Plant Mol. Biol. 59, 229–238 (2005).

34. M. Borg, F. Berger, Chromatin remodelling during male gametophyte development. Plant J. 83, 177–188 (2015).

35. R. Yelagandula, et al., The histone variant H2A.W defines heterochromatin and promotes chromatin condensation in Arabidopsis. Cell 158, 98–109 (2014).

36. M. D. Meneghini, M. Wu, H. D. Madhani, Conserved histone variant H2A.Z protects euchromatin from the ectopic spread of silent heterochromatin. Cell 112, 725–736 (2003).

37. B. P. M. Ngou, M. Wyler, M. W. Schmid, Y. Kadota, K. Shirasu, Evolutionary trajectory of pattern recognition receptors in plants. Nat. Commun. 15, 308 (2024).

38. R. L. López-Marqués, et al., Intracellular targeting signals and lipid specificity determinants of the ALA/ALIS P4-ATPase complex reside in the catalytic ALA α-subunit. Mol. Biol. Cell 21, 791–801 (2010).

39. S. C. McDowell, et al., Loss of the Arabidopsis thaliana P4-ATPases ALA6 and ALA7 impairs pollen fitness and alters the pollen tube plasma membrane. Front. Plant Sci. 6, 197 (2015).

40. G. Pilot, R. Pratelli, F. Gaymard, Y. Meyer, H. Sentenac, Five-group distribution of the Shaker-like K+ channel family in higher plants. J. Mol. Evol. 56, 418–434 (2003).

41. K. Mouline, et al., Pollen tube development and competitive ability are impaired by disruption of a Shaker K(+) channel in Arabidopsis. Genes Dev. 16, 339–350 (2002).

42. E. Michard, et al., Glutamate receptor-like genes form Ca2+ channels in pollen tubes and are regulated by pistil D-serine. Science 332, 434–437 (2011).

43. M. Kasahara, et al., An adenylyl cyclase with a phosphodiesterase domain in basal plants with a motile sperm system. Sci. Rep. 6, 39232 (2016).

44. C. Yamamoto, et al., The cAMP signaling module regulates sperm motility in the liverwort Marchantia polymorpha. Proc. Natl. Acad. Sci. U. S. A. 121, e2322211121 (2024).

45. Y. Liu, et al., The conserved plant sterility gene HAP2 functions after attachment of fusogenic membranes in Chlamydomonas and Plasmodium gametes. Genes Dev. 22, 1051–1068 (2008).

46. H. Yang, et al., Malectin/Malectin-like domain-containing proteins: A repertoire of cell surface molecules with broad functional potential. Cell Surf. 7, 100056 (2021).

47. Z. Ge, et al., Arabidopsis pollen tube integrity and sperm release are regulated by RALF-mediated signaling. Science 358, 1596–1600 (2017).

48. M. A. Mecchia, et al., The single Marchantia polymorpha FERONIA homolog reveals an ancestral role in regulating cellular expansion and integrity. Development 149 (2022).

49. J. M. Abercrombie, B. C. O’Meara, A. R. Moffatt, J. H. Williams, Developmental evolution of flowering plant pollen tube cell walls: callose synthase (CalS) gene expression patterns. Evodevo 2, 14 (2011).

50. J.-C. Mollet, C. Leroux, F. Dardelle, A. Lehner, Cell wall composition, biosynthesis and remodeling during pollen tube growth. Plants 2, 107–147 (2013).

51. K. Kobayashi, K. Awai, K.-I. Takamiya, H. Ohta, Arabidopsis type B monogalactosyldiacylglycerol synthase genes are expressed during pollen tube growth and induced by phosphate starvation. Plant Physiol. 134, 640–648 (2004).

52. R. Yatomi, S. Nakamura, N. Nakamura, Immunochemical and cytochemical detection of wall components of germinated pollen of gymnosperms. Grana 41, 21–28 (2002).

53. Y. Toyama, T. Higashiyama, Submicron-scale chromatin architecture in Cycas revoluta pollen nuclei. Mol. Reprod. Dev. 91, e23726 (2024).

54. L. Liu, T. Wang, Male gametophyte development in flowering plants: A story of quarantine and sacrifice. J. Plant Physiol. 258–259, 153365 (2021).

55. M. Hasebe, et al., Phylogeny of gymnosperms inferred fromrbcL gene sequences. Bot Mag Tokyo 105, 673–679 (1992).

56. S. M. Chaw, C. L. Parkinson, Y. Cheng, T. M. Vincent, J. D. Palmer, Seed plant phylogeny inferred from all three plant genomes: monophyly of extant gymnosperms and origin of Gnetales from conifers. Proc. Natl. Acad. Sci. U. S. A. 97, 4086–4091 (2000).

57. N. J. Wickett, et al., Phylotranscriptomic analysis of the origin and early diversification of land plants. Proc. Natl. Acad. Sci. U. S. A. 111, E4859–68 (2014).

58. Å. Åkerman, Über die Chemotaxis der Marchantia-Spermatozoiden. Z. Bot 2, 94–103 (1910).

59. C. J. Brokaw, Chemotaxis of Bracken Spermatozoids : The Role of Bimalate Ions. J. Exp. Biol. 35, 192–196 (1958).

60. B. J. Haas, et al., De novo transcript sequence reconstruction from RNA-seq using the Trinity platform for reference generation and analysis. Nat. Protoc. 8, 1494–1512 (2013).

61. N. L. Bray, H. Pimentel, P. Melsted, L. Pachter, Near-optimal probabilistic RNA-seq quantification. Nat. Biotechnol. 34, 525–527 (2016).

