## Supplemental information for "Intermediate Evolutionary State of Motile Sperm and Pollen Tubes in the Extant Gymnosperm *Cycas revoluta*"

**This PDF file includes:**

Figures S1 to S4
Tables S1 to S13
Legends for Movies S1 to S3
Legends for Dataset S1 to S2
SI References

**Other supporting materials for this manuscript include the following:**

Movies S1 to S3
Dataset S1 to S2

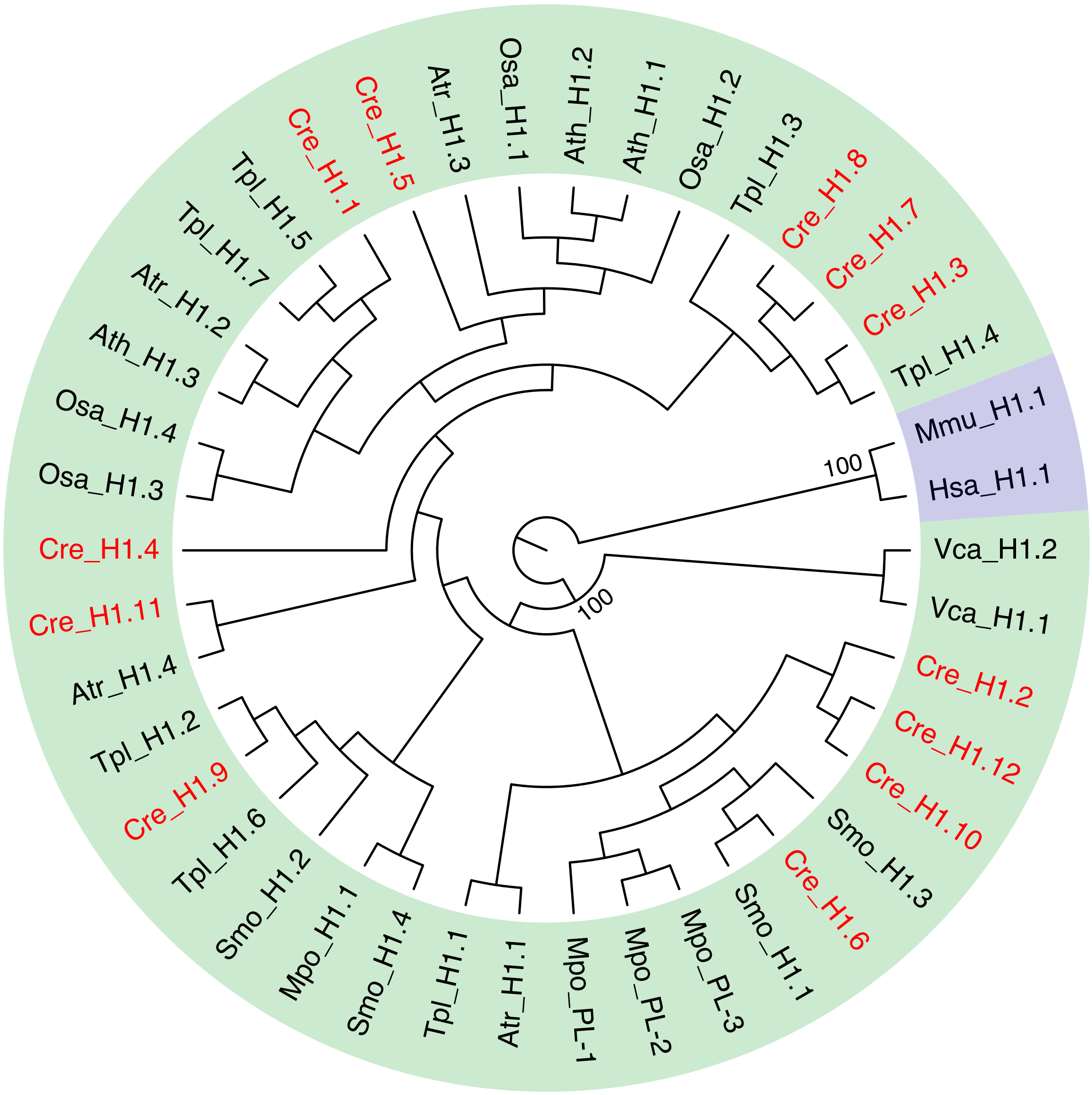


**Fig S1.** Maximum-likelihood phylogenetic tree constructed from the protein sequences of the H1 variant candidates of *C. revoluta* and the other H1 variants listed in Table S5 (1,000 bootstrap replicates). The tree was rooted using metazoan variants. Variants of land plants and green algae are shaded in green, and metazoan variants are shaded in blue. The variants shown in red text are of *C. revoluta*. Ath, *A. thaliana*; Atr, *A. trichopoda*; Cre, *C. revoluta*; Hmo, *H. sapiens*; Mmu, *M. musculus*; Mpo, *M. polymorpha*; Osa, *O. sativa*; Smo, *S. moellendorffii*; Tpl, *T. plicata*; Vca, *V. carteri*.

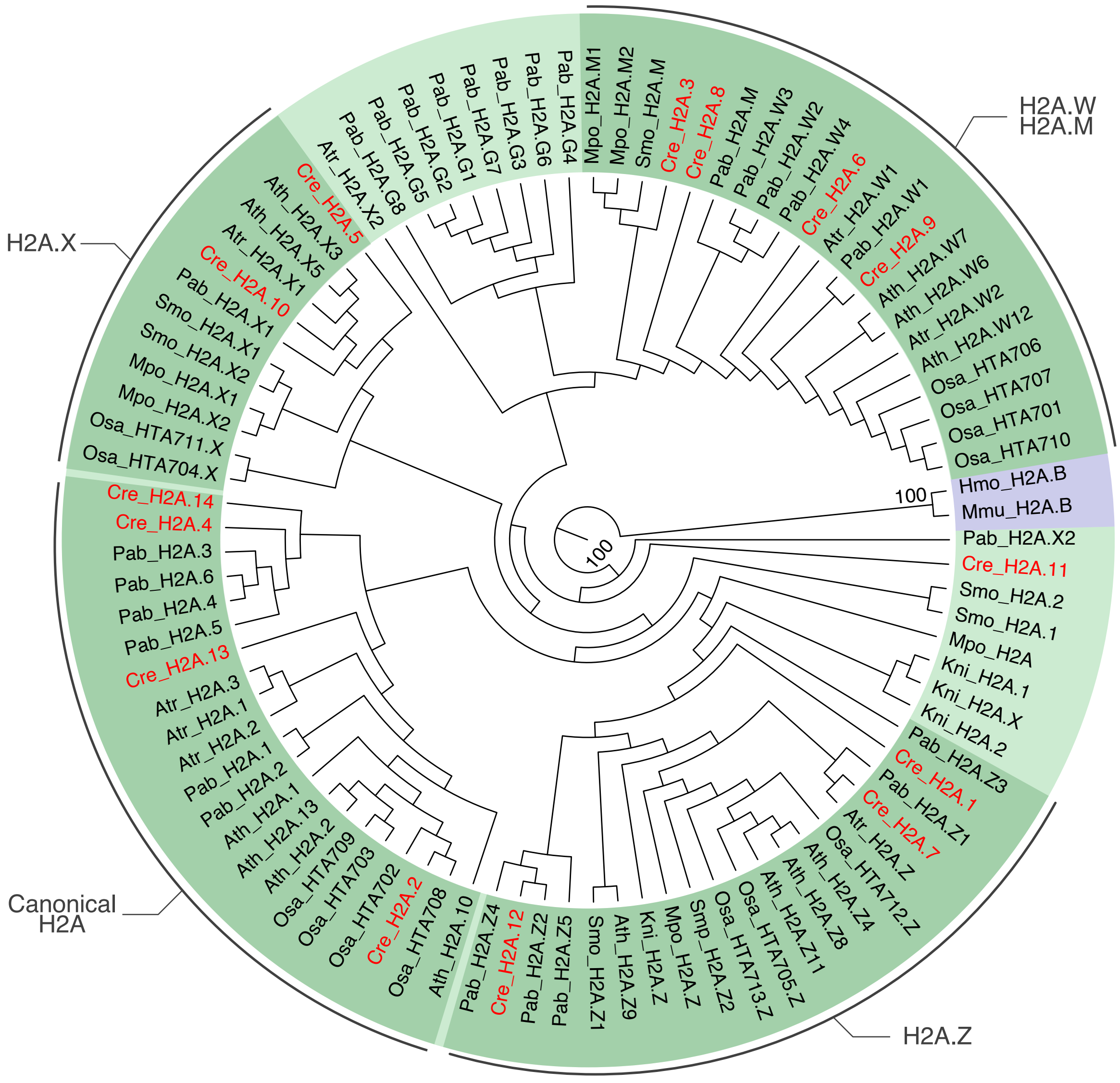


**Fig S2.** Maximum-likelihood phylogenetic tree constructed from the protein sequences of the H2A variant candidates of *C. revoluta* and the other H2A variants listed in Table S6 (1,000 bootstrap replicates). The tree was rooted using metazoan variants. Variants of land plants and green algae are shaded in green, and metazoan variants are shaded in blue. The variants shown in red text are of *C. revoluta*. H2A variant clades are highlighted, canonical H2A, H2A.Z, H2A.X, and H2A.W (H2A.M). *K. nitens*, Kni; *P. abies*, Pab.

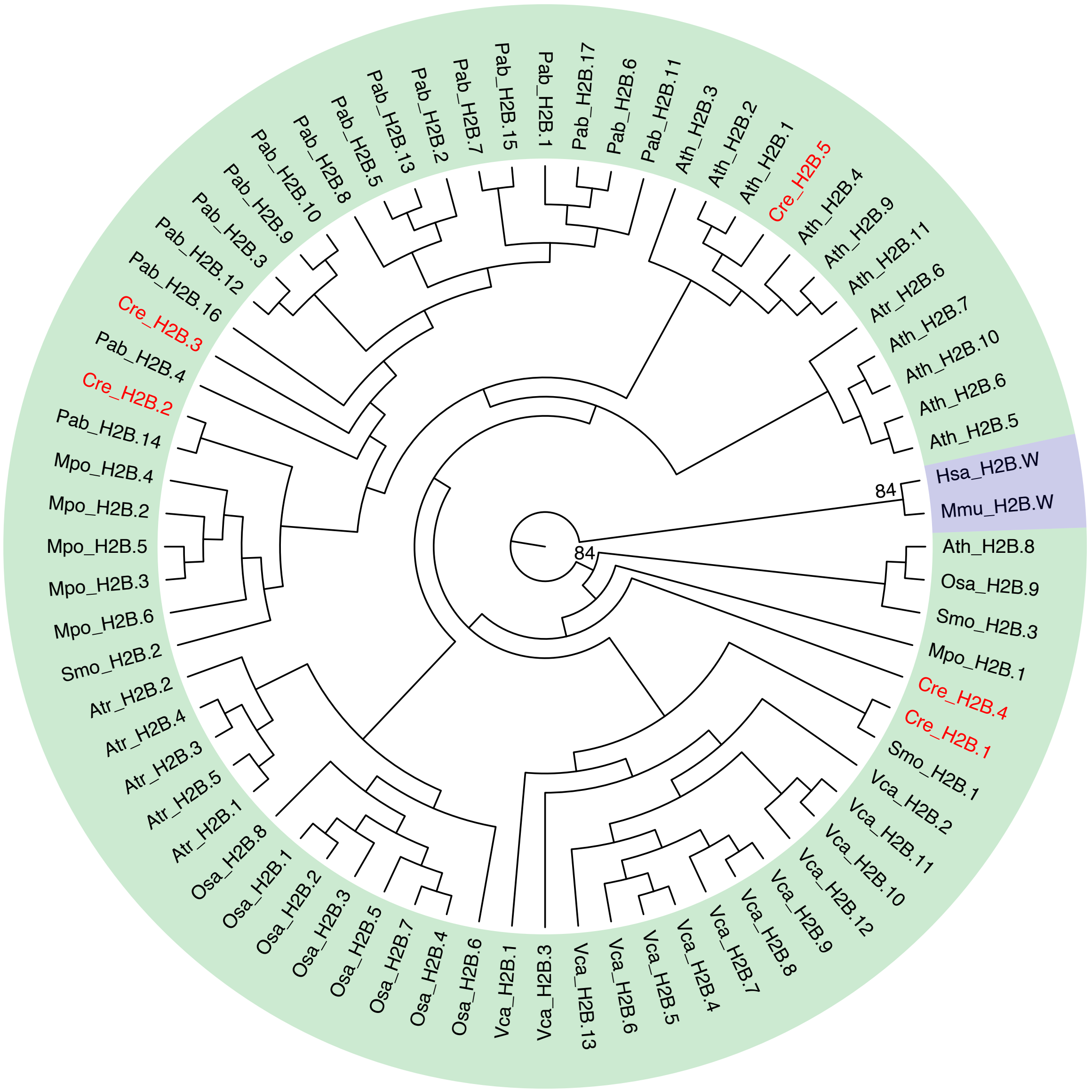


**Fig S3.** Maximum-likelihood phylogenetic tree constructed from the protein sequences of the H2B variant candidates of *C. revoluta* and the other H2B variants listed in Table S7 (1,000 bootstrap replicates). The tree was rooted using metazoan variants. Variants of land plants and green algae are shaded in green, and metazoan variants are shaded in blue. The variants shown in red text are of *C. revoluta*.

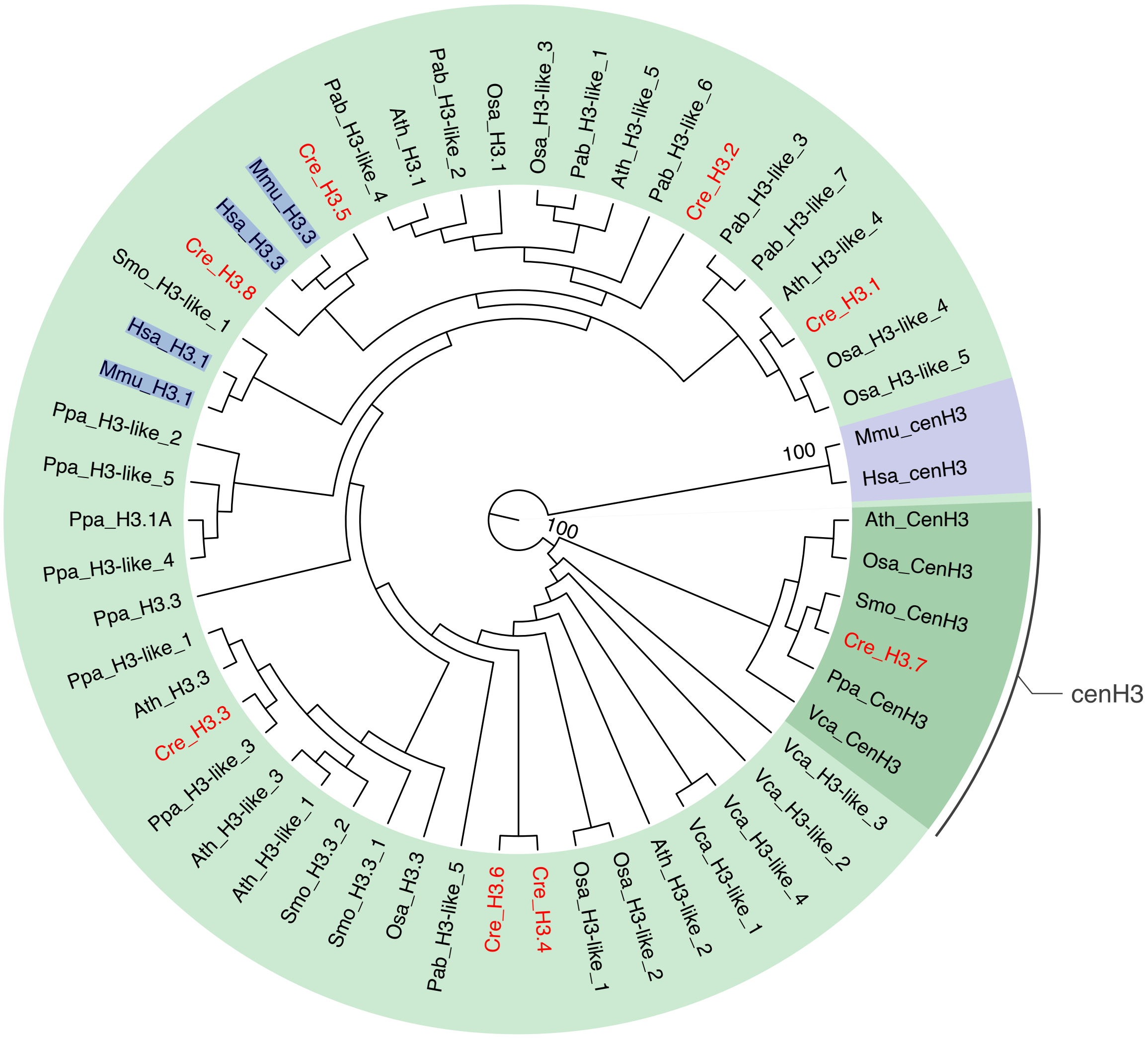


**Fig S4.** Maximum-likelihood phylogenetic tree constructed from the protein sequences of the H3 variant candidates of *C. revoluta* and the other H3 variants listed in Table S8 (1,000 bootstrap replicates). The tree was rooted using metazoan variants. Variants of land plants and green algae are shaded in green, and metazoan variants are shaded in blue. The variants shown in red text are of *C. revoluta*. H3 variant clade cenH3 is highlighted. *P. patens*, Ppa.

**Table S1.** BUSCO summary analyzed for 77,449 assembled CDSs.

| Datasets | Complete | | Fragmented | Missing |
|---|---|---|---|---|
| | (Single-copy) | (Duplicated) | | |
| eukaryota (N=255) | 92.5% | | 2.0% | 5.5% |
| | (48.2%) | (44.3%) | | |
| viridiplantae (N=425) | 90.8% | | 1.2% | 8.0% |
| | (53.4%) | (37.4%) | | |
| embryophyta (N=1614) | 84.0% | | 1.9% | 14.1% |
| | (51.9%) | (32.0%) | | |

**Table S2.** The number of all or tissue-specifically expressed CDSs for each tissue group.

| Tissue Groups | Number of CDSs | | |
|---|---|---|---|
| | TPM>0 | TPM>2 | SPM>0.90 |
| Leaf | 62,154 | 21,365 | 1,083 |
| Flower | 41,894 | 21,619 | 475 |
| Female | 72,142 | 34,826 | 3,926 |
| Male | 70,094 | 18,523 | 5,679 |

**Table S3.** The list of histone variant candidates of *C. revoluta*

| Histone Variant | Name in tree | CDS ID | Length (aa) | Conserved domain Accession | E-value | Isoforms |
|---|---|---|---|---|---|---|
| H1 | Cre_H1.1 | Cre11077_c0_g1_i1.p1 | 259 | smart00526 | 5.93005E-19 | 1 |
| | Cre_H1.2 | Cre11547_c0_g1_i1.p1 | 290 | smart00526 | 2.33203E-15 | 1 |
| | Cre_H1.3 | Cre162_c0_g2_i1.p1 | 301 | smart00526 | 3.06189E-11 | 2 |
| | Cre_H1.4 | Cre162367_c0_g1_i1.p1 | 311 | smart00526 | 4.77901E-20 | 1 |
| | Cre_H1.5 | Cre168854_c0_g1_i1.p1 | 272 | smart00526 | 1.76612E-15 | 1 |
| | Cre_H1.6 | Cre19923_c0_g4_i1.p1 | 180 | cl00073 | 1.34234E-06 | 1 |
| | Cre_H1.7 | Cre21779_c0_g1_i3.p1 | 210 | smart00526 | 1.32226E-17 | 2 |
| | Cre_H1.8 | Cre21779_c0_g2_i1.p1 | 196 | smart00526 | 3.79222E-19 | 1 |
| | Cre_H1.9 | Cre2382_c0_g1_i11.p1 | 463 | smart00526 | 1.03284E-18 | 1 |
| | Cre_H1.10 | Cre58080_c0_g1_i8.p1 | 193 | smart00526 | 2.45092E-16 | 1 |
| | Cre_H1.11 | Cre77243_c0_g1_i1.p1 | 211 | smart00526 | 2.76814E-18 | 1 |
| | Cre_H1.12 | Cre8675_c0_g2_i3.p1 | 310 | smart00526 | 1.3143E-12 | 2 |
| H2A | Cre_H2A.1 | Cre10136_c0_g1_i1.p1 | 143 | PLN00154 | 2.80488E-78 | 1 |
| | Cre_H2A.2 | Cre10846_c0_g1_i2.p1 | 139 | PLN00157 | 1.20962E-76 | 1 |
| | Cre_H2A.3 | Cre169384_c0_g1_i1.p1 | 140 | cl30549 | 1.19096E-20 | 1 |
| | Cre_H2A.4 | Cre177147_c0_g1_i1.p1 | 132 | PLN00157 | 4.79574E-71 | 1 |
| | Cre_H2A.5 | Cre185859_c0_g1_i1.p1 | 139 | cl29681 | 7.28445E-75 | 1 |
| | Cre_H2A.6 | Cre2187_c0_g1_i1.p1 | 145 | cl29679 | 4.11103E-64 | 2 |
| | Cre_H2A.7 | Cre2296_c0_g1_i1.p2 | 154 | PLN00154 | 2.74763E-77 | 1 |
| | Cre_H2A.8 | Cre30775_c0_g1_i1.p1 | 141 | cl30549 | 1.34379E-62 | 1 |
| | Cre_H2A.9 | Cre34931_c0_g1_i2.p1 | 143 | cl29679 | 1.48368E-66 | 1 |
| | Cre_H2A.10 | Cre3662_c0_g1_i1.p1 | 139 | PLN00156 | 4.2768E-83 | 1 |
| | Cre_H2A.11 | Cre403_c0_g1_i2.p1 | 142 | PTZ00017 | 6.67857E-72 | 4 |
| | Cre_H2A.12 | Cre5366_c0_g1_i1.p1 | 141 | PLN00154 | 6.73023E-75 | 1 |
| | Cre_H2A.13 | Cre64536_c0_g1_i1.p1 | 136 | PLN00157 | 5.57113E-71 | 1 |
| | Cre_H2A.14 | Cre78358_c0_g1_i2.p2 | 132 | PLN00157 | 1.72941E-71 | 1 |
| H2B | Cre_H2B.1 | Cre102769_c0_g1_i1.p1 | 130 | cd22910 | 2.80031E-58 | 1 |
| | Cre_H2B.2 | Cre538_c0_g1_i1.p1 | 150 | cd22910 | 1.5191E-58 | 5 |
| | Cre_H2B.3 | Cre538_c0_g2_i6.p1 | 143 | cd22910 | 2.31154E-59 | 5 |
| | Cre_H2B.4 | Cre5935_c0_g1_i1.p1 | 213 | cd22910 | 2.91301E-53 | 1 |
| | Cre_H2B.5 | Cre820_c0_g1_i24.p1 | 153 | cd22910 | 2.70146E-58 | 1 |
| H3 | Cre_H3.1 | Cre11950_c0_g2_i1.p2 | 138 | PTZ00018 | 2.3102E-75 | 1 |
| | Cre_H3.2 | Cre1437_c0_g1_i1.p1 | 137 | PTZ00018 | 6.067E-87 | 1 |
| | Cre_H3.3 | Cre1937_c0_g2_i2.p1 | 137 | PTZ00018 | 6.16709E-90 | 1 |
| | Cre_H3.4 | Cre1949_c0_g1_i1.p1 | 137 | PTZ00018 | 6.16709E-90 | 1 |
| | Cre_H3.5 | Cre2766_c0_g1_i1.p1 | 137 | PTZ00018 | 9.05502E-90 | 11 |
| | Cre_H3.6 | Cre2766_c0_g2_i1.p1 | 137 | PTZ00018 | 2.80932E-88 | 1 |
| | Cre_H3.7 | Cre4900_c0_g1_i1.p1 | 161 | cd22911 | 2.60205E-55 | 1 |
| | Cre_H3.8 | Cre8959_c0_g1_i5.p1 | 156 | PTZ00018 | 1.28729E-91 | 2 |

**Table S4.** All isoforms for the histone variant candidates of *C. revoluta* and their raw expression levels.

| Histone Variant | Name in Figure 3 | Expression value (TPM) | | | | | CDS ID |
|---|---|---|---|---|---|---|---|
| | | Sp | Po | Fe | Fl | Le | |
| H1 | Cre_H1.1 | 0.0 | 5.5 | 52.4 | 262.2 | 243.9 | Cre11077_c0_g1_i1.p1 |
| | Cre_H1.2 | 0.0 | 5.3 | 20.2 | 18.1 | 16.1 | Cre11547_c0_g1_i1.p1 |
| | Cre_H1.3.i1 | 0.0 | 1.3 | 15.0 | 68.8 | 46.5 | Cre162_c0_g2_i1.p1 |
| | Cre_H1.3.i2 | 0.0 | 0.9 | 0.1 | 42.4 | 1.9 | Cre162_c0_g2_i2.p1 |
| | Cre_H1.4 | 0.0 | 0.2 | 2.4 | 3.5 | 12.1 | Cre162367_c0_g1_i1.p1 |
| | Cre_H1.5 | 0.0 | 8.9 | 63.7 | 179.9 | 106.5 | Cre168854_c0_g1_i1.p1 |
| | Cre_H1.6 | 2.9 | 25.1 | 0.0 | 0.0 | 0.0 | Cre19923_c0_g4_i1.p1 |
| | Cre_H1.7.i1 | 3.0 | 2.9 | 0.1 | 0.0 | 0.0 | Cre21779_c0_g1_i3.p1 |
| | Cre_H1.7.i2 | 0.6 | 2.9 | 5.3 | 10.0 | 8.3 | Cre21779_c0_g1_i2.p1 |
| | Cre_H1.8 | 2.6 | 1.4 | 6.9 | 2.3 | 3.6 | Cre21779_c0_g2_i1.p1 |
| | Cre_H1.9 | 4.2 | 8.0 | 48.0 | 44.4 | 7.0 | Cre2382_c0_g1_i11.p1 |
| | Cre_H1.10 | 0.5 | 0.6 | 8.7 | 8.5 | 2.0 | Cre58080_c0_g1_i8.p1 |
| | Cre_H1.11 | 0.0 | 0.5 | 31.5 | 1.5 | 0.0 | Cre77243_c0_g1_i1.p1 |
| | Cre_H1.12.i1 | 0.2 | 5.3 | 15.1 | 0.0 | 6.4 | Cre8675_c0_g2_i3.p1 |
| | Cre_H1.12.i2 | 0.0 | 0.5 | 2.5 | 0.0 | 0.0 | Cre8675_c0_g2_i2.p2 |
| H2A | Cre_H2A.1 | 0.0 | 1.3 | 7.9 | 20.8 | 9.2 | Cre10136_c0_g1_i1.p1 |
| | Cre_H2A.2 | 0.0 | 4.1 | 22.6 | 41.3 | 205.7 | Cre10846_c0_g1_i2.p1 |
| | Cre_H2A.3 | 95.3 | 55.0 | 0.0 | 0.0 | 0.0 | Cre169384_c0_g1_i1.p1 |
| | Cre_H2A.4 | 0.0 | 1.7 | 45.1 | 152.4 | 91.6 | Cre177147_c0_g1_i1.p1 |
| | Cre_H2A.5 | 1.4 | 84.5 | 266.4 | 146.8 | 143.6 | Cre185859_c0_g1_i1.p1 |
| | Cre_H2A.6.i1 | 17.0 | 47.5 | 120.8 | 6.0 | 63.6 | Cre2187_c0_g1_i1.p1 |
| | Cre_H2A.6.i2 | 0.6 | 0.2 | 71.3 | 0.0 | 1.3 | Cre2187_c0_g1_i3.p2 |
| | Cre_H2A.7 | 0.0 | 18.4 | 41.6 | 17.3 | 10.3 | Cre2296_c0_g1_i1.p2 |
| | Cre_H2A.8 | 0.0 | 7.9 | 42.5 | 31.6 | 23.2 | Cre30775_c0_g1_i1.p1 |
| | Cre_H2A.9 | 0.0 | 0.0 | 3.7 | 3.9 | 0.6 | Cre34931_c0_g1_i2.p1 |
| | Cre_H2A.10 | 0.0 | 4.7 | 20.9 | 2.3 | 4.4 | Cre3662_c0_g1_i1.p1 |
| | Cre_H2A.11.i1 | 0.9 | 3.1 | 83.7 | 0.0 | 0.0 | Cre403_c0_g1_i2.p1 |
| | Cre_H2A.11.i2 | 17.0 | 33.7 | 13.3 | 0.0 | 0.0 | Cre403_c0_g1_i3.p1 |
| | Cre_H2A.11.i3 | 36.2 | 34.8 | 22.4 | 0.0 | 0.0 | Cre403_c0_g1_i5.p1 |
| | Cre_H2A.11.i4 | 9.2 | 12.4 | 8.6 | 0.0 | 0.0 | Cre403_c0_g1_i9.p1 |
| | Cre_H2A.12 | 0.7 | 9.1 | 11.5 | 5.1 | 10.3 | Cre5366_c0_g1_i1.p1 |
| | Cre_H2A.13 | 0.0 | 1.0 | 5.7 | 8.0 | 2.0 | Cre64536_c0_g1_i1.p1 |
| | Cre_H2A.14 | 29.2 | 21.9 | 87.1 | 51.3 | 70.3 | Cre78358_c0_g1_i2.p2 |
| H2B | Cre_H2B.1 | 9.7 | 16.4 | 10.5 | 0.0 | 0.0 | Cre102769_c0_g1_i1.p1 |
| | Cre_H2B.2.i1 | 0.0 | 3.6 | 5.8 | 0.0 | 6.6 | Cre538_c0_g1_i1.p1 |
| | Cre_H2B.2.i2 | 0.5 | 8.0 | 48.3 | 1096.5 | 577.7 | Cre538_c0_g1_i2.p1 |
| | Cre_H2B.2.i3 | 73.3 | 70.6 | 26.6 | 79.9 | 40.3 | Cre538_c0_g1_i3.p1 |
| | Cre_H2B.2.i4 | 0.5 | 10.7 | 62.8 | 90.6 | 72.1 | Cre538_c0_g1_i4.p1 |
| | Cre_H2B.2.i5 | 0.7 | 3.8 | 2.9 | 0.0 | 0.0 | Cre538_c0_g1_i5.p1 |
| | Cre_H2B.3.i1 | 0.8 | 6.2 | 11.6 | 18.8 | 16.8 | Cre538_c0_g2_i6.p1 |
| | Cre_H2B.3.i2 | 0.0 | 0.0 | 6.2 | 6.0 | 12.7 | Cre538_c0_g2_i1.p2 |
| | Cre_H2B.3.i3 | 0.0 | 2.5 | 51.4 | 0.0 | 0.0 | Cre538_c0_g2_i11.p1 |

| | | | | | | | |
|---|---|---|---|---|---|---|---|
| | Cre_H2B.3.i4 | 13.8 | 2.9 | 1.6 | 0.0 | 0.0 | Cre538_c0_g2_i5.p3 |
| | Cre_H2B.3.i5 | 0.1 | 6.4 | 0.0 | 0.0 | 0.0 | Cre538_c0_g2_i8.p2 |
| | Cre_H2B.4 | 0.0 | 2.2 | 3.6 | 0.0 | 0.1 | Cre5935_c0_g1_i1.p1 |
| | Cre_H2B.5.i1 | 9.0 | 7.6 | 6.3 | 46.6 | 24.0 | Cre820_c0_g1_i24.p1 |
| | Cre_H2B.5.i2 | 0.1 | 3.5 | 21.8 | 0.0 | 0.2 | Cre820_c0_g1_i26.p1 |
| H3 | Cre_H3.1 | 0.0 | 0.9 | 0.4 | 2.5 | 0.2 | Cre11950_c0_g2_i1.p2 |
| | Cre_H3.2 | 0.7 | 4.6 | 70.0 | 0.0 | 0.0 | Cre1437_c0_g1_i1.p1 |
| | Cre_H3.3 | 0.0 | 19.0 | 122.5 | 201.6 | 214.3 | Cre1937_c0_g2_i2.p1 |
| | Cre_H3.4 | 0.3 | 12.0 | 81.6 | 145.1 | 150.5 | Cre1949_c0_g1_i1.p1 |
| | Cre_H3.5.i1 | 2.1 | 7.2 | 6.2 | 0.0 | 0.0 | Cre2766_c0_g1_i1.p1 |
| | Cre_H3.5.i2 | 0.0 | 2.4 | 16.6 | 0.0 | 0.2 | Cre2766_c0_g1_i12.p1 |
| | Cre_H3.5.i3 | 18.4 | 18.4 | 14.8 | 0.0 | 0.0 | Cre2766_c0_g1_i13.p1 |
| | Cre_H3.5.i4 | 0.0 | 9.0 | 10.4 | 9.3 | 1.7 | Cre2766_c0_g1_i2.p2 |
| | Cre_H3.5.i5 | 1.1 | 1.7 | 3.3 | 0.0 | 0.0 | Cre2766_c0_g1_i3.p1 |
| | Cre_H3.5.i6 | 0.7 | 1.5 | 12.4 | 0.0 | 0.0 | Cre2766_c0_g1_i4.p1 |
| | Cre_H3.5.i7 | 0.0 | 2.4 | 0.0 | 9.3 | 0.5 | Cre2766_c0_g1_i5.p2 |
| | Cre_H3.5.i8 | 4.4 | 7.7 | 10.4 | 0.0 | 0.0 | Cre2766_c0_g1_i6.p1 |
| | Cre_H3.5.i9 | 0.7 | 1.5 | 12.4 | 0.0 | 0.0 | Cre2766_c0_g1_i7.p1 |
| | Cre_H3.5.i10 | 0.0 | 2.8 | 51.0 | 0.0 | 0.0 | Cre2766_c0_g1_i8.p1 |
| | Cre_H3.5.i11 | 1.9 | 5.6 | 15.5 | 0.0 | 0.0 | Cre2766_c0_g1_i9.p1 |
| | Cre_H3.6 | 56.0 | 64.7 | 33.9 | 71.1 | 297.5 | Cre2766_c0_g2_i1.p1 |
| | Cre_H3.7 | 7.5 | 9.2 | 3.1 | 1.5 | 1.9 | Cre4900_c0_g1_i1.p1 |
| | Cre_H3.8.i1 | 0.3 | 1.5 | 3.0 | 16.1 | 1.8 | Cre8959_c0_g1_i5.p1 |
| | Cre_H3.8.i2 | 14.3 | 18.7 | 14.1 | 0.0 | 0.9 | Cre8959_c0_g1_i3.p2 |
| | Cre_H3.8.i3 | 1.4 | 3.5 | 6.9 | 1.7 | 0.2 | Cre8959_c0_g1_i9.p1 |

*Tissues: Sperm (Sp), Pollen tube with Sperm (Po), Female (Fe), Flower (Fl), Leaf (Le).

**Table S5.** Accession numbers of the histone variants used for constructing H1 phylogenetic tree in Figure S1.

| Species | Reference Accession Number | Reference | Name in tree | Length (aa) |
|---|---|---|---|---|
| *Arabidopsis thaliana* | AT1G06760 | (1), TAIR | Ath_H1.1 | 274 |
| *Arabidopsis thaliana* | AT2G30620 | (1), TAIR | Ath_H1.2 | 273 |
| *Arabidopsis thaliana* | AT2G18050 | (1), TAIR | Ath_H1.3 | 167 |
| *Oryza sativa* | AK099211 | (2), NCBI | Osa_H1.1 | 293 |
| *Oryza sativa* | AK102642 | (2), NCBI | Osa_H1.2 | 278 |
| *Oryza sativa* | AK069863 | (2), NCBI | Osa_H1.3 | 240 |
| *Oryza sativa* | AK073668 | (2), NCBI | Osa_H1.4 | 188 |
| *Amborella trichopoda* | evm_27.TU.AmTr_v1.0_ scaffold00091.20 | Phytozome (Amborella trichopoda v1.0) | Atr_H1.1 | 199 |
| *Amborella trichopoda* | evm_27.TU.AmTr_v1.0_ scaffold00185.23 | Phytozome (Amborella trichopoda v1.0) | Atr_H1.2 | 207 |
| *Amborella trichopoda* | evm_27.TU.AmTr_v1.0_ scaffold00002.487 | Phytozome (Amborella trichopoda v1.0) | Atr_H1.3 | 243 |
| *Amborella trichopoda* | evm_27.TU.AmTr_v1.0_ scaffold00057.90 | Phytozome (Amborella trichopoda v1.0) | Atr_H1.4 | 191 |
| *Thuja plicata* | Thupl.29378171s0002 | Phytozome (Thuja plicata v3.1c) | Tpl_H1.1 | 161 |
| *Thuja plicata* | Thupl.29381632s0011 | Phytozome (Thuja plicata v3.1c) | Tpl_H1.2 | 379 |
| *Thuja plicata* | Thupl.29377942s0009 | Phytozome (Thuja plicata v3.1c) | Tpl_H1.3 | 246 |
| *Thuja plicata* | Thupl.29382418s0012 | Phytozome (Thuja plicata v3.1c) | Tpl_H1.4 | 287 |
| *Thuja plicata* | Thupl.29377851s0006 | Phytozome (Thuja plicata v3.1c) | Tpl_H1.5 | 229 |
| *Thuja plicata* | Thupl.29380178s0004 | Phytozome (Thuja plicata v3.1c) | Tpl_H1.6 | 255 |
| *Thuja plicata* | Thupl.29377851s0005 | Phytozome (Thuja plicata v3.1c) | Tpl_H1.7 | 250 |
| *Selaginella moellendorffii* | 423943 | Phytozome (Selaginella moellendorffii v1.0) | Smo_H1.1 | 113 |
| *Selaginella moellendorffii* | 438299 | Phytozome (Selaginella moellendorffii v1.0) | Smo_H1.2 | 237 |
| *Selaginella moellendorffii* | 27494 | Phytozome (Selaginella moellendorffii v1.0) | Smo_H1.3 | 149 |
| *Selaginella moellendorffii* | 440806 | Phytozome (Selaginella moellendorffii v1.0) | Smo_H1.4 | 378 |
| *Marchantia polymorpha* | MppBR5_0015s0990 | MarpolBase (Mp Reference Genome v7.1) | Mpo_H1.1 | 299 |
| *Marchantia polymorpha* | BAU71552 | NCBI | Mpo_PL-1 | 251 |
| *Marchantia polymorpha* | PTQ35141 | NCBI | Mpo_PL-2 | 344 |
| *Marchantia polymorpha* | PTQ32223 | NCBI | Mpo_PL-3 | 306 |
| *Volvox carteri* | Q08864 | (3), NCBI | Vca_H1.1 | 261 |
| *Volvox carteri* | Q08865 | (3), NCBI | Vca_H1.2 | 241 |
| *Homo sapiens* | X57130 | (3), NCBI | Hsa_H1.1 | 215 |
| *Mus musculus* | S43949 | (3), Uniprot | Mmu_H1.1 | 213 |

**Table S6.** Accession numbers of the histone variants used for constructing H2A phylogenetic tree in Figure S2.

| Species | Reference Accession Number | Reference | Name in tree | Length (aa) |
|---|---|---|---|---|
| *Arabidopsis thaliana* | AT1G08880 | (4), TAIR | Ath_H2A.X5 | 142 |
| *Arabidopsis thaliana* | AT1G51060 | (4), TAIR | Ath_H2A.10 | 132 |
| *Arabidopsis thaliana* | AT1G52740 | (4), TAIR | Ath_H2A.Z9 | 134 |
| *Arabidopsis thaliana* | AT1G54690 | (4), TAIR | Ath_H2A.X3 | 142 |
| *Arabidopsis thaliana* | AT2G38810 | (4), TAIR | Ath_H2A.Z8 | 136 |
| *Arabidopsis thaliana* | AT3G20670 | (4), TAIR | Ath_H2A.13 | 132 |
| *Arabidopsis thaliana* | AT3G54560 | (4), TAIR | Ath_H2A.Z11 | 136 |
| *Arabidopsis thaliana* | AT4G13570 | (4), TAIR | Ath_H2A.Z4 | 124 |
| *Arabidopsis thaliana* | AT4G27230 | (4), TAIR | Ath_H2A.2 | 131 |
| *Arabidopsis thaliana* | AT5G02560 | (4), TAIR | Ath_H2A.W12 | 153 |
| *Arabidopsis thaliana* | AT5G27670 | (4), TAIR | Ath_H2A.W7 | 150 |
| *Arabidopsis thaliana* | AT5G54640 | (4), TAIR | Ath_H2A.1 | 130 |
| *Arabidopsis thaliana* | AT5G59870 | (4), TAIR | Ath_H2A.W6 | 150 |
| *Oryza satiiva* | NP_001411163 | (2), NCBI | Osa_HTA702 | 134 |
| *Oryza satiiva* | AK071511 | (2), NCBI | Osa_HTA703 | 135 |
| *Oryza satiiva* | AK059228 | (2), NCBI | Osa_HTA708 | 135 |
| *Oryza satiiva* | AK099888 | (2), NCBI | Osa_HTA709 | 135 |
| *Oryza satiiva* | AK064299 | (2), NCBI | Osa_HTA704.X | 137 |
| *Oryza satiiva* | AK121752 | (2), NCBI | Osa_HTA711.X | 138 |
| *Oryza satiiva* | NP_001415043 | (2), NCBI | Osa_HTA701 | 159 |
| *Oryza satiiva* | AK067406 | (2), NCBI | Osa_HTA706 | 163 |
| *Oryza satiiva* | AK074018 | (2), NCBI | Osa_HTA707 | 156 |
| *Oryza satiiva* | AK121750 | (2), NCBI | Osa_HTA710 | 159 |
| *Oryza satiiva* | AK121515 | (2), NCBI | Osa_HTA705.Z | 139 |
| *Oryza satiiva* | AK121533 | (2), NCBI | Osa_HTA712.Z | 138 |
| *Oryza satiiva* | AK120299 | (2), NCBI | Osa_HTA713.Z | 137 |
| *Amborella trichopoda* | evm_27.TU.AmTr_v1.0_ scaffold00057.272 | (4), Phytozome (Amborella trichopoda v1.0) | Atr_H2A.W1 | 147 |
| *Amborella trichopoda* | evm_27.TU.AmTr_v1.0_ scaffold00168.14 | (4), Phytozome (Amborella trichopoda v1.0) | Atr_H2A.W2 | 147 |
| *Amborella trichopoda* | evm_27.TU.AmTr_v1.0_ scaffold00197.10 | (4), Phytozome (Amborella trichopoda v1.0) | Atr_H2A.X1 | 140 |
| *Amborella trichopoda* | evm_27.TU.AmTr_v1.0_ scaffold00055.171 | (4), Phytozome (Amborella trichopoda v1.0) | Atr_H2A.1 | 134 |
| *Amborella trichopoda* | evm_27.TU.AmTr_v1.0_ scaffold00115.21 | (4), Phytozome (Amborella trichopoda v1.0) | Atr_H2A.2 | 134 |
| *Amborella trichopoda* | evm_27.TU.AmTr_v1.0_ scaffold00058.136 | (4), Phytozome (Amborella trichopoda v1.0) | Atr_H2A.X2 | 137 |
| *Amborella trichopoda* | evm_27.TU.AmTr_v1.0_ scaffold00016.224 | (4), Phytozome (Amborella trichopoda v1.0) | Atr_H2A.3 | 126 |
| *Amborella trichopoda* | evm_27.TU.AmTr_v1.0_ scaffold00007.104 | (4), Phytozome (Amborella trichopoda v1.0) | Atr_H2A.Z | 99 |
| *Picea abies* | MA_10434390g0020 | (4), Diurnal plant tools | Pab_H2A.3 | 129 |
| *Picea abies* | MA_191819g0010 | (4), Diurnal plant tools | Pab_H2A.5 | 131 |
| *Picea abies* | MA_94328g0020 | (4), Diurnal plant tools | Pab_H2A.X1 | 138 |
| *Picea abies* | MA_22203g0020 | (4), Diurnal plant tools | Pab_H2A.2 | 140 |
| *Picea abies* | MA_427046g0010 | (4), Diurnal plant tools | Pab_H2A.1 | 140 |
| *Picea abies* | MA_9379473g0010 | (4), Diurnal plant tools | Pab_H2A.X2 | 145 |
| *Picea abies* | MA_10425800g0010 | (4), Diurnal plant tools | Pab_H2A.G2 | 144 |
| *Picea abies* | MA_74667g0010 | (4), Diurnal plant tools | Pab_H2A.G1 | 144 |
| *Picea abies* | MA_169054g0020 | (4), Diurnal plant tools | Pab_H2A.G8 | 147 |

| | | | | |
|---|---|---|---|---|
| *Picea abies* | MA_11514g0010 | (4), Diurnal plant tools | Pab_H2A.G6 | 145 |
| *Picea abies* | MA_213048g0010 | (4), Diurnal plant tools | Pab_H2A.G5 | 143 |
| *Picea abies* | MA_865576g0010 | (4), Diurnal plant tools | Pab_H2A.G4 | 144 |
| *Picea abies* | MA_10426943g0010 | (4), Diurnal plant tools | Pab_H2A.G3 | 144 |
| *Picea abies* | MA_924620g0010 | (4), Diurnal plant tools | Pab_H2A.W3 | 141 |
| *Picea abies* | MA_962778g0010 | (4), Diurnal plant tools | Pab_H2A.W2 | 140 |
| *Picea abies* | MA_100910g0010 | (4), Diurnal plant tools | Pab_H2A.M | 135 |
| *Picea abies* | MA_7320190g0010 | (4), Diurnal plant tools | Pab_H2A.W1 | 141 |
| *Picea abies* | MA_93628g0020 | (4), Diurnal plant tools | Pab_H2A.W4 | 142 |
| *Picea abies* | MA_3822944g0010 | (4), Diurnal plant tools | Pab_H2A.G7 | 131 |
| *Picea abies* | MA_114554g0010 | (4), Diurnal plant tools | Pab_H2A.Z3 | 142 |
| *Picea abies* | MA_100300g0010 | (4), Diurnal plant tools | Pab_H2A.Z1 | 145 |
| *Picea abies* | MA_68993g0010 | (4), Diurnal plant tools | Pab_H2A.Z4 | 124 |
| *Picea abies* | MA_19162g0010 | (4), Diurnal plant tools | Pab_H2A.Z5 | 145 |
| *Picea abies* | MA_323386g0010 | (4), Diurnal plant tools | Pab_H2A.Z2 | 116 |
| *Picea abies* | MA_562932g0010 | (4), Diurnal plant tools | Pab_H2A.4 | 133 |
| *Picea abies* | MA_10434390g0010 | (4), Diurnal plant tools | Pab_H2A.6 | 134 |
| *Selaginella moellendorffii* | 73637 | (4), Phytozome (Selaginella moellendorffii V1.0) | Smo_H2A.X1 | 140 |
| *Selaginella moellendorffii* | 170948 | (4), Phytozome (Selaginella moellendorffii V1.0) | Smo_H2A.X2 | 136 |
| *Selaginella moellendorffii* | 38507 | (4), Phytozome (Selaginella moellendorffii V1.0) | Smo_H2A.M | 136 |
| *Selaginella moellendorffii* | 38585 | (4), Phytozome (Selaginella moellendorffii V1.0) | Smo_H2A.Z1 | 139 |
| *Selaginella moellendorffii* | 76653 | (4), Phytozome (Selaginella moellendorffii V1.0) | Smo_H2A.1 | 132 |
| *Selaginella moellendorffii* | 166626 | (4), Phytozome (Selaginella moellendorffii V1.0) | Smo_H2A.2 | 137 |
| *Selaginella moellendorffii* | 122533 | (4), Phytozome (Selaginella moellendorffii V1.0) | Smp_H2A.Z2 | 135 |
| *Marchantia polymorpha* | Mp5g19440 | (4), MarpolBase | Mpo_H2A.M1 | 160 |
| *Marchantia polymorpha* | Mp3g00840 | (4), MarpolBase | Mpo_H2A.M2 | 147 |
| *Marchantia polymorpha* | Mp1g09020 | (4), MarpolBase | Mpo_H2A.Z | 141 |
| *Marchantia polymorpha* | Mp2g00760 | (4), MarpolBase | Mpo_H2A.X1 | 139 |
| *Marchantia polymorpha* | Mp4g21750 | (4), MarpolBase | Mpo_H2A.X2 | 138 |
| *Marchantia polymorpha* | Mp3g02370 | (4), MarpolBase | Mpo_H2A | 129 |
| *Klebsormidium nitens* | kfl00106_0030_v1.1 | (4), NIES-2285 V1.1 Transcripts (Predicted Protein) | Kni_H2A.1 | 174 |
| *Klebsormidium nitens* | kfl00107_0200_v1.1 | (4), NIES-2285 V1.1 Transcripts (Predicted Protein) | Kni_H2A.Z | 136 |
| *Klebsormidium nitens* | kfl00125_0190_v1.1 | (4), NIES-2285 V1.1 Transcripts (Predicted Protein) | Kni_H2A.X | 147 |
| *Klebsormidium nitens* | kfl00099_g21_v1.1 | (4), NIES-2285 V1.1 Transcripts (Predicted Protein) | Kni_H2A.2 | 135 |
| *Homo sapiens* | P0C5Y9 | (5), Uniprot | Hmo_H2A.B | 115 |
| *Mus musculus* | S4R1M3 | (5), Uniprot | Mmu_H2A.B | 115 |

**Table S7.** Accession numbers of the histone variants used for constructing H2B phylogenetic tree in Figure S3.

| Species | Reference Accession Number | Reference | Name in tree | Length (aa) |
|---|---|---|---|---|
| *Arabidopsis thaliana* | AT1G07790 | (1), TAIR | Ath_H2B.1 | 148 |
| *Arabidopsis thaliana* | AT5G22880 | (1), TAIR | Ath_H2B.2 | 145 |
| *Arabidopsis thaliana* | AT2G28720 | (1), TAIR | Ath_H2B.3 | 151 |
| *Arabidopsis thaliana* | AT5G59910 | (1), TAIR | Ath_H2B.4 | 150 |
| *Arabidopsis thaliana* | AT2G37470 | (1), TAIR | Ath_H2B.5 | 138 |
| *Arabidopsis thaliana* | AT3G53650 | (1), TAIR | Ath_H2B.6 | 138 |
| *Arabidopsis thaliana* | AT3G09480 | (1), TAIR | Ath_H2B.7 | 126 |
| *Arabidopsis thaliana* | AT1G08170 | (1), TAIR | Ath_H2B.8 | 243 |
| *Arabidopsis thaliana* | AT3G45980 | (1), TAIR | Ath_H2B.9 | 150 |
| *Arabidopsis thaliana* | AT5G02570 | (1), TAIR | Ath_H2B.10 | 132 |
| *Arabidopsis thaliana* | AT3G46030 | (1), TAIR | Ath_H2B.11 | 145 |
| *Oryza sativa* | LOC_Os01g05610 | (6), Phytozome | Osa_H2B.1 | 153 |
| *Oryza sativa* | LOC_Os01g05630 | (6), Phytozome | Osa_H2B.2 | 153 |
| *Oryza sativa* | LOC_Os01g05900 | (6), Phytozome | Osa_H2B.3 | 153 |
| *Oryza sativa* | LOC_Os01g05970 | (6), Phytozome | Osa_H2B.4 | 153 |
| *Oryza sativa* | LOC_Os01g06010 | (6), Phytozome | Osa_H2B.5 | 155 |
| *Oryza sativa* | LOC_Os01g62230 | (6), Phytozome | Osa_H2B.6 | 139 |
| *Oryza sativa* | LOC_Os05g49860 | (6), Phytozome | Osa_H2B.7 | 152 |
| *Oryza sativa* | LOC_Os08g38300 | (6), Phytozome | Osa_H2B.8 | 150 |
| *Oryza sativa* | LOC_Os09g39730 | (6), Phytozome | Osa_H2B.9 | 194 |
| *Amborella trichopoda* | evm_27.TU.AmTr_v1.0_ scaffold00010.258 | (6), Phytozome | Atr_H2B.1 | 150 |
| *Amborella trichopoda* | evm_27.TU.AmTr_v1.0_ scaffold00012.26 | (6), Phytozome | Atr_H2B.2 | 148 |
| *Amborella trichopoda* | evm_27.TU.AmTr_v1.0_ scaffold00012.38 | (6), Phytozome | Atr_H2B.3 | 139 |
| *Amborella trichopoda* | evm_27.TU.AmTr_v1.0_ scaffold00012.39 | (6), Phytozome | Atr_H2B.4 | 148 |
| *Amborella trichopoda* | evm_27.TU.AmTr_v1.0_ scaffold00012.42 | (6), Phytozome | Atr_H2B.5 | 139 |
| *Amborella trichopoda* | evm_27.TU.AmTr_v1.0_ scaffold00085.91 | (6), Phytozome | Atr_H2B.6 | 136 |
| *Picea abies* | lcl\|MA_10428006g0030 | (6), Congenie | Pab_H2B.1 | 141 |
| *Picea abies* | lcl\|MA_138049g0010 | (6), Congenie | Pab_H2B.2 | 138 |
| *Picea abies* | lcl\|MA_8065g0010 | (6), Congenie | Pab_H2B.3 | 142 |
| *Picea abies* | lcl\|MA_111473g0010 | (6), Congenie | Pab_H2B.4 | 139 |
| *Picea abies* | lcl\|MA_10238123g0010 | (6), Congenie | Pab_H2B.5 | 143 |
| *Picea abies* | lcl\|MA_10435143g0010 | (6), Congenie | Pab_H2B.6 | 141 |
| *Picea abies* | lcl\|MA_77198g0010 | (6), Congenie | Pab_H2B.7 | 162 |
| *Picea abies* | lcl\|MA_10434433g0010 | (6), Congenie | Pab_H2B.8 | 185 |
| *Picea abies* | lcl\|MA_10436516g0010 | (6), Congenie | Pab_H2B.9 | 141 |
| *Picea abies* | lcl\|MA_10436516g0020 | (6), Congenie | Pab_H2B.10 | 141 |
| *Picea abies* | lcl\|MA_38565g0010 | (6), Congenie | Pab_H2B.11 | 141 |
| *Picea abies* | lcl\|MA_454575g0010 | (6), Congenie | Pab_H2B.12 | 142 |

| | | | | |
|---|---|---|---|---|
| *Picea abies* | lcllMA_1011737g0010 | (6), Congenie | Pab_H2B.13 | 143 |
| *Picea abies* | lcllMA_9756g0010 | (6), Congenie | Pab_H2B.14 | 157 |
| *Picea abies* | lcllMA_6078786g0010 | (6), Congenie | Pab_H2B.15 | 164 |
| *Picea abies* | lcllMA_17795g0010 | (6), Congenie | Pab_H2B.16 | 140 |
| *Picea abies* | lcllMA_9293473g0010 | (6), Congenie | Pab_H2B.17 | 141 |
| *Selaginella moellendorffii* | 182308 | (6), Phytozome | Smo_H2B.1 | 141 |
| *Selaginella moellendorffii* | 73644 | (6), Phytozome | Smo_H2B.2 | 146 |
| *Selaginella moellendorffii* | 76616 | (6), Phytozome | Smo_H2B.3 | 121 |
| *Marchantia polymorpha* | Mp1g27660.1 | (6), MarpolBase | Mpo_H2B.1 | 185 |
| *Marchantia polymorpha* | Mp6g04070.1 | (6), MarpolBase | Mpo_H2B.2 | 142 |
| *Marchantia polymorpha* | Mp6g07010.1 | (6), MarpolBase | Mpo_H2B.3 | 139 |
| *Marchantia polymorpha* | Mp7g18140.1 | (6), MarpolBase | Mpo_H2B.4 | 142 |
| *Marchantia polymorpha* | Mp4g13450.1 | (6), MarpolBase | Mpo_H2B.5 | 145 |
| *Marchantia polymorpha* | Mp7g05430.1 | (6), MarpolBase | Mpo_H2B.6 | 141 |
| *Volvox carteri* | Vocar.0001s0460 | (6), MarpolBase | Vca_H2B.1 | 121 |
| *Volvox carteri* | Vocar.0001s0881 | (6), Phytozome | Vca_H2B.2 | 156 |
| *Volvox carteri* | Vocar.0002s0359 | (6), Phytozome | Vca_H2B.3 | 160 |
| *Volvox carteri* | Vocar.0002s0389 | (6), Phytozome | Vca_H2B.4 | 161 |
| *Volvox carteri* | Vocar.0002s0392 | (6), Phytozome | Vca_H2B.5 | 159 |
| *Volvox carteri* | Vocar.0005s0053 | (6), Phytozome | Vca_H2B.6 | 156 |
| *Volvox carteri* | Vocar.0006s0197 | (6), Phytozome | Vca_H2B.7 | 154 |
| *Volvox carteri* | Vocar.0016s0272 | (6), Phytozome | Vca_H2B.8 | 155 |
| *Volvox carteri* | Vocar.0016s0281 | (6), Phytozome | Vca_H2B.9 | 154 |
| *Volvox carteri* | Vocar.0027s0139 | (6), Phytozome | Vca_H2B.10 | 155 |
| *Volvox carteri* | Vocar.0027s0143 | (6), Phytozome | Vca_H2B.11 | 155 |
| *Volvox carteri* | Vocar.0027s0148 | (6), Phytozome | Vca_H2B.12 | 155 |
| *Volvox carteri* | Vocar.0028s0099 | (6), Phytozome | Vca_H2B.13 | 157 |
| *Homo sapiens* | Q7Z2G1 | (5), Uniprot | Hsa_H2B.W | 153 |
| *Mus musculus* | Q9DAB5 | (5), Uniprot | Mmu_H2B.W | 224 |

**Table S8.** Accession numbers of the histone variants used for constructing H3 phylogenetic tree in Figure S4.

| Species | Reference Accession Number | Reference | Name in tree | Length (aa) |
|---|---|---|---|---|
| *Arabidopsis thaliana* | AT1G01370 | (7), TAIR | Ath_CenH3 | 179 |
| *Arabidopsis thaliana* | AT1G09200 | (7), TAIR | Ath_H3.1 | 137 |
| *Arabidopsis thaliana* | AT4G40030 | (7), TAIR | Ath_H3.3 | 165 |
| *Arabidopsis thaliana* | AT1G13370 | (7), TAIR | Ath_H3-like_1 | 137 |
| *Arabidopsis thaliana* | AT1G19890 | (7), TAIR | Ath_H3-like_2 | 138 |
| *Arabidopsis thaliana* | AT1G75600 | (7), TAIR | Ath_H3-like_3 | 137 |
| *Arabidopsis thaliana* | AT5G12910 | (7), TAIR | Ath_H3-like_4 | 132 |
| *Arabidopsis thaliana* | AT5G65350 | (7), TAIR | Ath_H3-like_5 | 140 |
| *Oryza sativa* | Os05g41080 | (7), NCBI | Osa_CenH3 | 166 |
| *Oryza sativa* | Os01g64640 | (7), NCBI | Osa_H3.1 | 137 |
| *Oryza sativa* | Os03g27310 | (7), NCBI | Osa_H3.3 | 137 |
| *Oryza sativa* | Os02g25910 | (7), NCBI | Osa_H3-like_1 | 212 |
| *Oryza sativa* | Os02g25940 | (7), NCBI | Osa_H3-like_2 | 116 |
| *Oryza sativa* | Os06g06480 | (7), NCBI | Osa_H3-like_3 | 251 |
| *Oryza sativa* | Os12g22650 | (7), NCBI | Osa_H3-like_4 | 137 |
| *Oryza sativa* | Os12g22680 | (7), NCBI | Osa_H3-like_5 | 137 |
| *Picea abies* | MA_10432805g0020 | (7), Diurnal plant tools | Pab_H3-like_1 | 137 |
| *Picea abies* | MA_113838g0010 | (7), Diurnal plant tools | Pab_H3-like_2 | 137 |
| *Picea abies* | MA_197719g0010 | (7), Diurnal plant tools | Pab_H3-like_3 | 137 |
| *Picea abies* | MA_210354g0010 | (7), Diurnal plant tools | Pab_H3-like_4 | 137 |
| *Picea abies* | MA_259842g0010 | (7), Diurnal plant tools | Pab_H3-like_5 | 137 |
| *Picea abies* | MA_475294g0010 | (7), Diurnal plant tools | Pab_H3-like_6 | 137 |
| *Picea abies* | MA_56411g0010 | (7), Diurnal plant tools | Pab_H3-like_7 | 137 |
| *Seraginella moellendorffii* | 171192 | (7), Phytozome (Selaginella moellendorffii V1.0) | Smo_CenH3 | 127 |
| *Seraginella moellendorffii* | 176603 | (7), Phytozome (Selaginella moellendorffii V1.0) | Smo_H3.3_1 | 137 |
| *Seraginella moellendorffii* | 269004 | (7), Phytozome (Selaginella moellendorffii V1.0) | Smo_H3.3_2 | 137 |
| *Seraginella moellendorffii* | 103546 | (7), Phytozome (Selaginella moellendorffii V1.0) | Smo_H3-like_1 | 137 |
| *Physcomitrella patens* | Pp1s568 | (7), Phytozome (Physcomitrium patens V3.3) | Ppa_CenH3 | 129 |
| *Physcomitrella patens* | Pp1s12 | (7), Phytozome (Physcomitrium patens V3.3) | Ppa_H3.1A | 137 |
| *Physcomitrella patens* | Pp1s3_368V6 | (7), Phytozome (Physcomitrium patens V3.3) | Ppa_H3.3 | 144 |
| *Physcomitrella patens* | Pp1s1963_1V6 | (7), Phytozome (Physcomitrium patens V3.3) | Ppa_H3-like_1 | 137 |
| *Physcomitrella patens* | Pp1s26_96V6 | (7), Phytozome (Physcomitrium patens V3.3) | Ppa_H3-like_2 | 139 |
| *Physcomitrella patens* | Pp1s3_594V6 | (7), Phytozome (Physcomitrium patens V3.3) | Ppa_H3-like_3 | 137 |

| | | | | |
|---|---|---|---|---|
| *Physcomitrella patens* | Pp1s35_164V6 | (7), Phytozome (Physcomitrium patens V3.3) | Ppa_H3-like_4 | 137 |
| *Physcomitrella patens* | Pp1s9_383V6 | (7), Phytozome (Physcomitrium patens V3.3) | Ppa_H3-like_5 | 162 |
| *Volvox carteri* | Vocar20005857m | (7), Phytozome (Volvox carteri v2.1) | Vca_CenH3 | 144 |
| *Volvox carteri* | Vocar20002453m | (7), Phytozome (Volvox carteri v2.1) | Vca_H3-like_1 | 136 |
| *Volvox carteri* | Vocar20003601m | (7), Phytozome (Volvox carteri v2.1) | Vca_H3-like_2 | 136 |
| *Volvox carteri* | Vocar20005447m | (7), Phytozome (Volvox carteri v2.1) | Vca_H3-like_3 | 174 |
| *Volvox carteri* | Vocar20010992m | (7), Phytozome (Volvox carteri v2.1) | Vca_H3-like_4 | 136 |
| *Homo sapiens* | P68431 | (5), Uniprot | Hsa_H3.1 | 136 |
| *Homo sapiens* | P84243 | (5), Uniprot | Hsa_H3.3 | 136 |
| *Homo sapiens* | P49450-1 | (5), Uniprot | Hsa_cenH3 | 140 |
| *Mus musculus* | P68433 | (5), Uniprot | Mmu_H3.1 | 136 |
| *Mus musculus* | P84244 | (5), Uniprot | Mmu_H3.3 | 136 |
| *Mus musculus* | O35216 | (5), Uniprot | Mmu_cenH3 | 134 |

**Table S9.** List of the genes predicted to localize sperm or sperm cell plasma membrane indicated in Figure 4B.

| Species | Reference Accession Number | Description (Egg Nog Mapper) | Gene Type | Gene Name | Homologus Gene (Tair) |
|---|---|---|---|---|---|
| *A. thaliana* | AT2G07040 | LRR receptor-like serine threonine-protein kinase RLK | RLKs | LRR-RLK | - |
| *A. thaliana* | AT4G18640 | belongs to the protein kinase superfamily | RLKs | LRR-RLK | - |
| *A. thaliana* | AT4G20790 | Leucine-rich repeat | RLKs | LRR-RLK | - |
| *A. thaliana* | AT4G28670 | cysteine-rich receptor-like protein kinase 43 | RLKs | DUF26-RLK | - |
| *A. thaliana* | AT4G34440 | Proline-rich receptor-like protein kinase PERK5 | RLKs | PERK | - |
| *A. thaliana* | AT5G45840 | Leucine rich repeat N-terminal domain | RLKs | MDIS1 | - |
| *A. thaliana* | AT1G09930 | oligopeptide transporter | Transporter | OPT2 | - |
| *A. thaliana* | AT4G32500 | Potassium channel | Transporter | AKT5 | - |
| *A. thaliana* | AT5G55930 | oligopeptide transporter | Transporter | OPT1 | - |
| *A. thaliana* | AT1G79450 | ALA-Interacting Subunit | Other Membrane Protein | CDC50 | - |
| *Z. mays* | Zm00001e000952_P001 | Protein kinase domain | RLKs | PERK | AT4G34440 |
| *Z. mays* | Zm00001e002559_P001 | Protein kinase domain | RLKs | PERK | AT2G48170 |
| *Z. mays* | Zm00001e003582_P002 | Protein kinase domain | RLKs | LRR-RLK | AT4G31250 |
| *Z. mays* | Zm00001e010334_P002 | Protein tyrosine kinase | RLKs | LRR-RLK | AT1G78980 |
| *Z. mays* | Zm00001e013764_P002 | Protein tyrosine kinase | RLKs | LRR-RLK | AT4G31250 |
| *Z. mays* | Zm00001e021908_P001 | Protein kinase domain | RLKs | LRR-RLK | AT5G35390 |
| *Z. mays* | Zm00001e025126_P001 | Protein tyrosine kinase | RLKs | LRR-RLK | AT4G31250 |
| *Z. mays* | Zm00001e027954_P001 | Protein kinase domain | RLKs | PERK | AT1G23540 |
| *Z. mays* | Zm00001e029870_P002 | Carbohydrate-binding protein of the ER | RLKs | M/MLD-RLK | AT2G21480 |
| *Z. mays* | Zm00001e030728_P002 | Protein kinase domain | RLKs | PERK | AT1G23540 |
| *Z. mays* | Zm00001e036240_P001 | Leucine rich repeat N-terminal domain | LRR-RLK | LRR-RLK | AT3G42880 |
| *Z. mays* | Zm00001e036642_P001 | Carbohydrate-binding protein of the ER | M/MLD-RLK | M/MLD-RLK | AT2G21480 |
| *Z. mays* | Zm00001e015126_P001 | Lung seven transmembrane receptor | Other Receptor | 7TMs | AT5G18520 |
| *Z. mays* | Zm00001e041205_P001 | Lung seven transmembrane receptor | Other Receptor | 7TMs | AT5G18520 |
| *Z. mays* | Zm00001e027976_P001 | KHA, dimerisation domain of potassium ion channel | Transporter | AKT1 | AT2G26650 |

| | | | | | |
|---|---|---|---|---|---|
| *Z. mays* | Zm00001e032178_P002 | LEM3 (ligand-effect modulator 3) family / CDC50 family | Other Membrane Protein | CDC50 | AT3G12740 |
| *A. trichopoda* | AMTR_s00016p00259140 | receptor-like protein kinase | RLKs | M/MLD-RLK | AT3G51550 |
| *A. trichopoda* | AMTR_s00024p00049240 | RECEPTOR-like protein kinase | RLKs | M/MLD-RLK | AT4G39110 |
| *A. trichopoda* | AMTR_s00024p00252010 | inactive leucine-rich repeat receptor-like protein kinase | RLKs | LRR-RLK | AT3G42880 |
| *A. trichopoda* | AMTR_s00025p00206890 | LRR receptor-like serine threonine-protein kinase | RLKs | LRR-RLK | AT3G20190 |
| *A. trichopoda* | AMTR_s00025p00216900 | strubbelig-receptor family | RLKs | LRR-RLK | AT1G53730 |
| *A. trichopoda* | AMTR_s00045p00071050 | RECEPTOR-like protein kinase | RLKs | M/MLD-RLK | AT4G39110 |
| *A. trichopoda* | AMTR_s00059p00070500 | belongs to the protein kinase superfamily | RLKs | PERK | AT2G48170 |
| *A. trichopoda* | AMTR_s00059p00211320 | Protein kinase domain | RLKs | LRR-RLK | AT4G23740 |
| *A. trichopoda* | AMTR_s00085p00158310 | Cysteine-rich receptor-like protein kinase | RLKs | DUF26-RLK | AT1G70520 |
| *A. trichopoda* | AMTR_s00130p00117400 | LRR receptor-like serine threonine-protein kinase | RLKs | LRR-RLK | AT5G45840 |
| *A. trichopoda* | AMTR_s00039p00168460 | Lung seven transmembrane receptor | Other Receptor | 7TMs | AT5G18520 |
| *A. trichopoda* | AMTR_s00020p00030660 | Cytochrome b561 and DOMON domain-containing protein | Transporter | CRR | AT3G25290 |
| *A. trichopoda* | AMTR_s00061p00190940 | oligopeptide transporter | Transporter | OPT1 | AT5G55930 |
| *A. trichopoda* | AMTR_s00044p00055920 | ALA-Interacting Subunit | Other Membrane Protein | CDC50 | AT3G12740 |
| *C. revoluta* | Cre5347_c0_g1_i1.p1 | G-type lectin S-receptor-like serine threonine-protein kinase SD2-5 | RLKs | G-lectin-RLK | AT4G32300 |
| *C. revoluta* | Cre4183_c0_g1_i12.p2 | Glutamate-gated receptor that probably acts as non-selective cation channel | Other Receptor | GLR | AT1G42540 |
| *C. revoluta* | Cre4484_c0_g1_i4.p1 | Potassium channel | Transporter | AKT1 | AT2G26650 |
| *C. revoluta* | Cre13827_c0_g4_i1.p1 | 3',5'-cyclic-nucleotide phosphodiesterase activity | Other Membrane Protein | CAPE | Mp7g08500 |
| *C. revoluta* | Cre799_c0_g1_i2.p1 | ALA-interacting subunit | Other Membrane Protein | CDC50 | AT1G16360 |
| *C. revoluta* | Cre17458_c0_g1_i2.p1 | oligopeptide transporter *TPM < 2 in sperm | Transporter | OPT1 | AT5G55930 |
| *M. polumorpha* | Mp3g24110.1 | Lectin-domain containing receptor kinase | RLKs | L-lectin-RLK | AT3G16530 |
| *M. polumorpha* | Mp4g18760.1 | Belongs to the protein kinase superfamily | RLKs | LRR-RLK | AT1G56145 |

**Table S10.** Putative plasma membrane-localized genes among cycad sperm-specific CDSs that were not annotated as signal transduction with the COG classification.

| Gene ID | Blast Description | Gene type |
|---|---|---|
| Cre14452_c0_g1_i5.p1 | Gamete Expressed 2 (GEX2) | Gamete Fusion |
| Cre19556_c0_g1_i38.p1 | Hapless 2 / Generative Cell Specific 1 | Gamete Fusion |
| Cre36147_c0_g1_i1.p1 | LRR receptor-like serine/threonine-protein kinase GSO1 | RLK |
| Cre2140_c0_g1_i4.p1 | Leucine-rich repeat protein kinase family protein | RLK |
| Cre1022_c0_g3_i4.p1 | H[+]-ATPase 11 | Transporter |
| Cre17458_c0_g1_i2.p1 | Oligopeptide transporter 1 (OTP1) | Transporter |
| Cre200646_c0_g1_i1.p1 | WAT1-related protein | Transporter |
| Cre2815_c0_g1_i1.p1 | PIN7 | Transporter |
| Cre31634_c0_g1_i1.p1 | inositol transporter 2 | Transporter |
| Cre3913_c0_g1_i5.p1 | autoinhibited Ca(2+)-ATPase 10 | Transporter |
| Cre4713_c0_g1_i2.p1 | inositol transporter 2 | Transporter |
| Cre51_c0_g1_i3.p1 | magnesium transporter 2 | Transporter |
| Cre5136_c0_g1_i69.p1 | sulfate transporter 3;1 | Transporter |
| Cre4889_c0_g1_i1.p1 | WAT1-related protein | Transporter |
| Cre5309_c0_g1_i1.p1 | Sugar transport protein 7 (STP) | Transporter |
| Cre5813_c0_g3_i2.p1 | zinc transporter 1 precursor | Transporter |
| Cre879_c0_g2_i1.p1 | sugar transporter 1 (STP) | Transporter |
| Cre32595_c0_g1_i41.p1 | GABA transporter 1 | Transporter |
| Cre1881_c0_g1_i16.p1 | Choline transporter protein 1 | Transporter |
| Cre8441_c0_g1_i4.p1 | Cation-chloride cotransporter 1 | Transporter |
| Cre32986_c0_g1_i2.p1 | Voltage-gated potassium channels | ion channel |
| Cre4089_c0_g1_i4.p1 | Mechanosensitive ion channel family protein | ion channel |
| Cre828_c0_g1_i3.p1 | MLO-like protein | ion channel |
| Cre7682_c0_g1_i3.p1 | MLO-like protein | ion channel |
| Cre19556_c0_g1_i38.p1 | O-Glycosyl hydrolases family 17 protein | GPI anchor |
| Cre2140_c0_g1_i4.p1 | O-Glycosyl hydrolases family 17 protein | GPI anchor |

**Table S11.** The raw expression levels of five candidate CDSs with malectin or malectin-like domains of *C. revoluta* for Figure 4A.

| CDS ID | Name in tree | Expression value (TPM) | | | | | E-Value | Conserved Domain Accession | Length (aa) |
|---|---|---|---|---|---|---|---|---|---|
| | | Sp | Po | Fe | Fl | Le | | | |
| Cre10439_c0_g1_i3.p1 | CreMal_1 | 0.0 | 0.3 | 16.7 | 0.2 | 3.0 | 1.5E-45 | pfam11721 | 676 |
| Cre124804_c0_g1_i1.p1 | CreMal_2 | 0.1 | 1.2 | 4.1 | 0.4 | 1.4 | 3.4E-88 | pfam12819 | 948 |
| Cre168901_c0_g1_i1.p1 | CreMal_3 | 0.1 | 1.0 | 4.7 | 0.9 | 3.2 | 1.4E-28 | cl38375 | 877 |
| Cre19238_c0_g1_i2.p1 | CreMal_4 | 0.0 | 0.4 | 2.3 | 0.3 | 0.6 | 1.1E-99 | pfam12819 | 946 |
| Cre3579_c0_g1_i2.p1 | CreMal_5 | 0.0 | 0.0 | 0.0 | 0.0 | 0.5 | 2.2E-60 | pfam12819 | 995 |

*Tissues: Sperm (Sp), Pollen tube with Sperm (Po), Female (Fe), Flower (Fl), Leaf (Le).

**Table S12.** Accession numbers of MD/MLD containing genes used for constructing the phylogenetic tree in Figure 4B.

| Type | Species | Accession Number | Name in tree | Length (aa) |
|---|---|---|---|---|
| MLD-RLK | *Arabidopsis thaliana* | AT1G30570 | AT1G30570_MLD-RLK_HERK2 | 850 |
| | *Arabidopsis thaliana* | AT2G21480 | AT2G21480_MLD-RLK_BUPS2 | 872 |
| | *Arabidopsis thaliana* | AT2G23200 | AT2G23200_MLD-RLK | 835 |
| | *Arabidopsis thaliana* | AT2G39360 | AT2G39360_MLD-RLK_CVY1 | 817 |
| | *Arabidopsis thaliana* | AT3G04690 | AT3G04690_MLD-RLK_ANX1 | 851 |
| | *Arabidopsis thaliana* | AT3G46290 | AT3G46290_MLD-RLK_HERK1 | 831 |
| | *Arabidopsis thaliana* | AT3G51550 | AT3G51550_MLD-RLK_FER | 896 |
| | *Arabidopsis thaliana* | AT4G00300 | AT4G00300_MLD-RLK | 479 |
| | *Arabidopsis thaliana* | AT4G39110.1 | AT4G39110.1_MLD-RLK_BUPS1 | 879 |
| | *Arabidopsis thaliana* | AT5G24010 | AT5G24010_MLD-RLK | 1704 |
| | *Arabidopsis thaliana* | AT5G28680 | AT5G28680_MLD-RLK_ANX2 | 859 |
| | *Arabidopsis thaliana* | AT5G38990 | AT5G38990_MLD-RLK_MDS1 | 881 |
| | *Arabidopsis thaliana* | AT5G39000 | AT5G39000_MLD-RLK_MDS2 | 874 |
| | *Arabidopsis thaliana* | AT5G39020 | AT5G39020_MLD-RLK_MDS3 | 814 |
| | *Arabidopsis thaliana* | AT5G39030 | AT5G39030_MLD-RLK_MDS4 | 807 |
| | *Arabidopsis thaliana* | AT5G54380 | AT5G54380_MLD-RLK_THE1 | 856 |
| | *Arabidopsis thaliana* | AT5G59700 | AT5G59700_MLD-RLK | 830 |
| | *Arabidopsis thaliana* | AT5G61350 | AT5G61350_MLD-RLK_ERU_CAP1 | 843 |
| | *Marchantia polymorpha* | MP4G15890 | MP4G15890_MpFER | 894 |
| MLD-LRR-RLK | *Arabidopsis thaliana* | AT1G05700 | AT1G05700_MLD-LRR-RLK | 884 |
| | *Arabidopsis thaliana* | AT1G07550 | AT1G07550_MLD-LRR-RLK | 865 |
| MLD-RLP | *Arabidopsis thaliana* | AT1G24485 | AT1G24485_MLD-RLP | 575 |
| | *Arabidopsis thaliana* | AT1G25570 | AT1G25570_MLD-RLP | 629 |
| LRR-MD-RLK | *Arabidopsis thaliana* | AT1G07650 | AT1G07650_LRR-MD-RLK | 1021 |
| | *Arabidopsis thaliana* | AT1G29720 | AT1G29720_LRR-MD-RLK | 1020 |
| M_Kinesin_like | *Arabidopsis thaliana* | AT1G72250 | AT1G72250_M_Kinesin_like | 1216 |
| | *Arabidopsis thaliana* | AT2G22610 | AT2G22610_M_Kinesin_like | 1084 |

**Table S13.** Male tissue (pollen tube) specifically expressed genes associated with cell wall / membrane / envelope biogenesis annotated with COG classification.

| Species | Reference Accession number | Description (Egg Nog Mapper) | PFAMs (Egg Nog Mapper) | Blast Anotation |
|---|---|---|---|---|
| *A. thaliana* | AT2G29040 | Exostosin family | Exostosin | GT11 |
| | AT3G07620 | Exostosin family | Exostosin | - |
| | AT4G16745 | Exostosin family | Exostosin | - |
| | AT5G11130 | Exostosin family | Exostosin | - |
| | AT5G25310 | Exostosin family | Exostosin | - |
| | AT2G17010 | mechanosensitive ion channel | MS_channel | MSL8 |
| | AT4G00234 | Mechanosensitive ion channel protein | MS_channel | - |
| | AT2G13680 | callose synthase | FKS1_dom1,Glucan_synthase,Vta1 | CALS5 |
| | AT2G11810 | Monogalactosyldiacylglycerol (MGDG) synthase | Glyco_tran_28_C,MGDG_synth | AtMGD3 |
| | AT5G20410 | Monogalactosyldiacylglycerol synthase 2 | Glyco_tran_28_C,MGDG_synth | AtMGD2 |
| | AT1G02000 | 4-epimerase | Epimerase,GDP_Man_Dehyd | GAE2 |
| | AT1G04920 | Belongs to the glycosyltransferase 1 family | Glycos_transf_1,S6PP,Sucrose_synth | - |
| | AT1G31070 | UDP-N-acetylglucosamine | UDPGP | - |
| | AT1G63180 | Bifunctional UDP-glucose 4-epimerase and UDP-xylose 4-epimerase | GDP_Man_Dehyd | - |
| | AT1G71697 | choline kinase | Choline_kin_N,Choline_kinase | - |
| | AT2G19690 | phospholipase | - | - |
| | AT2G19880 | Glycosyl transferase family 21 | Glyco_transf_21 | - |
| | AT3G46440 | udp-glucuronic acid decarboxylase | GDP_Man_Dehyd | - |
| | AT3G55590 | Mannose-1-phosphate guanylyltransferase | Hexapep,NTP_transferase | - |
| | AT4G07960 | xyloglucan glycosyltransferase 12 | Glyco_tranf_2_3,Glyco_trans_2_3 | - |
| | AT4G20100 | vacuolar amino acid transporter | PQ-loop | - |
| | AT4G29460 | phospholipase | - | - |
| | AT4G29470 | phospholipase | - | - |
| | AT4G37420 | Glycosyl transferase family 2 | Glyco_transf_92 | - |
| | AT5G03760 | Glycosyltransferase like family 2 | Glyco_tranf_2_3,Glyco_trans_2_3 | - |
| | AT5G11110 | Belongs to the glycosyltransferase 1 family | Glycos_transf_1,S6PP,Sucrose_synth | - |
| | AT5G12390 | Component of the peroxisomal and mitochondrial division machineries. | Fis1_TPR_C,Fis1_TPR_N | - |
| | AT5G40720 | Glycosyl transferase family 2 | Glyco_transf_92 | - |
| *Z. mays* | Zm00001e002306_P001 | Exostosin family | Exostosin | - |
| | Zm00001e004646_P001 | Exostosin family | Exostosin | AT2G20370 (MUR3) |

| | | | | |
|---|---|---|---|---|
| | Zm00001e020358_P006 | Exostosin family | Exostosin | - |
| | Zm00001e029766_P001 | 1,3-beta-glucan synthase component | FKS1_dom1,Glucan_synthase,Vta1 | AT2G13680 (CALS5) |
| | Zm00001e029772_P001 | Callose synthase 3-like | FKS1_dom1,Glucan_synthase,Vta1 | AT2G13680 (CALS5) |
| | Zm00001e016149_P001 | Monogalactosyldiacylglycerol (MGDG) synthase | Glyco_tran_28_C,MGDG_synth | AT5G20410 (AtMGD2) |
| | Zm00001e030744_P003 | GDP-mannose 4,6 dehydratase | Epimerase,GDP_Man_Dehyd | AT4G30440 (GAE1) |
| | Zm00001e036291_P001 | GDP-mannose 4,6 dehydratase | Epimerase,GDP_Man_Dehyd | AT4G30440 (GAE1) |
| | Zm00001e005818_P002 | Glycosyl transferase family 2 | Glyco_tranf_2_3,Glyco_trans_2_3 | - |
| | Zm00001e013001_P001 | rRNA small subunit methyltransferase G | CMAS | - |
| | Zm00001e023715_P001 | UDP-glucuronate 4-epimerase | - | - |
| | Zm00001e024185_P001 | galactosyl transferase GMA12/MNN10 family | Glyco_transf_34 | - |
| | Zm00001e025195_P001 | Belongs to the SIS family. GutQ KpsF subfamily | CBS,SIS | - |
| | Zm00001e032241_P002 | Choline/ethanolamine kinase | Choline_kin_N,Choline_kinase | - |
| *A. trichopoda* | AMTR_s00012p00237500 | xyloglucan galactosyltransferase | Exostosin | AT2G20370 (MUR3) |
| | AMTR_s00133p00039770 | Xyloglucan galactosyltransferase KATAMARI1 | Exostosin | - |
| | AMTR_s00001p00171160 | mechanosensitive ion channel | MS_channel | AT2G17010 (MSL8) |
| | AMTR_s00037p00187160 | callose synthase | FKS1_dom1,Glucan_synthase | - |
| | AMTR_s00150p00030620 | callose synthase | FKS1_dom1,Glucan_synthase,Vta1 | AT2G13680 (CALS5) |
| | AMTR_s00033p00201820 | Monogalactosyldiacylglycerol synthase | Glyco_tran_28_C,MGDG_synth | AT5G20410 (AtMGD2) |
| | AMTR_s00110p00097630 | UDP-glucuronate 4-epimerase | Epimerase,GDP_Man_Dehyd | AT1G02000 (GAE2) |
| | AMTR_s00059p00197160 | WD repeat-containing protein | WD40 | - |
| | AMTR_s00061p00145220 | Belongs to the glycosyltransferase 1 family | Glycos_transf_1,S6PP,Sucrose_synth | - |
| | AMTR_s00092p00147500 | udp-glucuronic acid decarboxylase | GDP_Man_Dehyd | - |
| *C. revoluta* | Cre39208_c0_g5_i2.p1 | Glycosyltransferase | Exostosin | AT5G11610 (Exostosin) |
| | Cre4089_c0_g1_i4.p1 | Mechanosensitive ion channel | MS_channel | AT2G17010 (MSL8) |
| | Cre6589_c1_g1_i3.p1 | Encoded by | Amino_oxidase,CMAS,NAD_binding_8 | - |

**Movie S1 (separate file).** The process of two sperm exiting from a pollen tube. The movie corresponds to the time-lapse image of Figure 1C.

**Movie S2 (separate file).** Isolated intact sperm swimming freely outside the pollen tubes. The movie corresponds to the merged images of Figure 1D.

**Movie S3 (separate file).** Isolated intact sperm swimming around neck cells as if attracted to them. The movie corresponds to the Figure 1E.

**Dataset S1 (separate file).** Nucleotide sequences of 77,449 CDSs.

**Dataset S2 (separate file).** Expression levels (TPM) of all CDSs in each sample.

### References


1. P. B. Talbert, *et al.*, A unified phylogeny-based nomenclature for histone variants. *Epigenetics Chromatin* **5**, 7 (2012).
2. Y. Hu, Y. Lai, Identification and expression analysis of rice histone genes. *Plant Physiol. Biochem.* **86**, 55–65 (2015).
3. H. E. Kasinsky, J. D. Lewis, J. B. Dacks, J. Ausió, Origin of H1 linker histones. *FASEB J.* **15**, 34–42 (2001).
4. T. Kawashima, *et al.*, Diversification of histone H2A variants during plant evolution. *Trends Plant Sci.* **20**, 419–425 (2015).
5. S. El Kennani, *et al.*, MS_HistoneDB, a manually curated resource for proteomic analysis of human and mouse histones. *Epigenetics Chromatin* **10**, 2 (2017).
6. D. Jiang, *et al.*, The evolution and functional divergence of the histone H2B family in plants. *PLoS Genet.* **16**, e1008964 (2020).
7. J. Cui, *et al.*, Genome-wide identification, evolutionary, and expression analyses of histone H3 variants in plants. *Biomed Res. Int.* **2015**, 341598 (2015).
